# Optimal Transport Method-Based Gene Filter (GF) Denoising Algorithm for Enhancing Spatially Resolved Transcriptomics Data

**DOI:** 10.1101/2023.07.01.547049

**Authors:** Lin Du, Jingmin Kang, Haixi Sun, Bohan Zhang

## Abstract

The recent advancements in spatially resolved transcriptomics (SRT) technology have enabled the acquisition of gene expression data at near- or sub-single-cell resolution, along with simultaneous imaging of physical locations. Nevertheless, necessary experimental procedures such as tissue fixation, permeabilization, and tissue removal inevitably induce the diffusion of transcribed molecules. Consequently, this leads to the partial capture of ex-situ transcripts in SRT data, thereby introducing a considerable amount of noise into the dataset. To address this issue, in this study, we focused on evaluating the diffusion pattern of individual genes within tissue regions and quantitatively calculating their signal-to-noise ratio (SNR). Through this analysis, we successfully identified “invalid genes” exhibiting widespread expression across tissue regions. Then by filtering out these genes, we effectively reduced the high noise level present in SRT data. To achieve this, we developed the gene filter denoising (GF) algorithm, which utilizes the optimal transport method to compute the gene diffusion coefficient and generate denoised SRT data. One notable advantage of our GF algorithm is its ability to fully “respect” the raw sequencing data, thereby avoiding the introduction of false positives often associated with traditional interpolation and modification denoising methods. Furthermore, we conducted comprehensive validation of GF, and the GF-denoised SRT data demonstrated substantial improvements in clustering, identification of differentially expressed genes (DEGs), and cell type annotation. Taken together, we believe that the GF denoising technique will serve as an essential and crucial step in exploring SRT data and investigating the underlying biological processes.

## Introduction

The spatially resolved transcriptomics (SRT) technology combines high-throughput gene sequencing with histology techniques to provide comprehensive gene expression data of the entire transcriptome, accompanied by spatially matched images such as Hematoxylin-eosin (H&E), immunofluorescence (IF), cell wall Calcofluor white, and nucleus 4,6-Diamidino-2-phenyl indole dihydrochloride (DAPI) staining^1^. One of the highlights of SRT data comprises the ability to integrate spatial location to analyze cell types, assisting in cell function mining, cell interaction, cell development, and other analyses. The experimental steps of SRT involve tissue section imaging and RNA sequencing. Ideally, each sequencing unit (referred to as a bead or spot in different technologies) at a specific location should capture only the transcripts released by cells located above it. However, due to the random diffusion of transcripts in the liquid experimental environment, SRT data tends to capture only a partial representation of ex-situ transcripts^2, 3^. This random diffusion introduces complex and severe noise issues in SRT data, which go beyond the drop-outs^4^ phenomena typically observed in single-cell RNA sequencing (scRNA-seq)^2, 5^.

There are three primary factors contributing to transcript diffusion. Firstly, during the process of tissue sectioning, frozen embedded tissue sections are affixed to the surface of the sequencing chip for tissue fixation and permeabilization. The sequencing chip beneath the tissue captures mRNA released through cell lysis, enabling the acquisition of positional information regarding gene expression. However, incomplete cell lysis and low quantities of released transcripts can lead to inadequate utilization of primers corresponding to the spots on the chip beneath the cells. Consequently, these spots may capture drifting mRNA from the surrounding environment^6, 7^. Conversely, excessive cell lysis and an abundance of released transcripts can cause an overflow of mRNA, which may drift to neighboring spots and be captured there. As the cell lysis process occurs in a liquid environment, mRNA drift is a random event devoid of direction, making precise control challenging. Secondly, transcript diffusion can occur during the reverse transcription and amplification stages of RNA sequencing. During reverse transcription, RNA needs to combine with reverse transcriptase to form a complex, and the structures of the enzymes or reaction conditions may result in the damage or leakage of RNA within the complex. Additionally, in cDNA synthesis, RNA needs to form a complex with primers, and incomplete binding or improper reaction conditions can cause primers to induce RNA diffusion^8^. Lastly, the utilization of inappropriate algorithms, parameters, or quality control standards during SRT data processing can also introduce noise, leading to an increased false positive rate in the SRT data^9^. Additionally, the occurrence of transcript swapping between tissues and backgrounds is extensively discussed in the SpotClean^7^ software article, providing a quantitative assessment of this phenomenon. In addition to addressing the well-known drop-outs phenomenon, the Sprod^5^ paper provides a thorough validation of the presence of complex spatial noise in SRT data, specifically examining the 10X Visium ovarian cancer dataset.

Numerous methods have been used to mitigate the noise present in SRT data. Initially, researchers turned to expression imputation techniques originally developed to tackle the drop-outs phenomenon in scRNA-seq^10^ data, such as MAGIC^11^, scImpute^12^, and SAVER^13^. However, these imputation methods only offered modest improvements and proved insufficient in effectively addressing the spatially correlated noise evident in SRT data^9^. As a result, subsequent efforts focused on developing algorithms tailored specifically for the complex noise patterns observed in SRT data. One such method is the Sprod^5^ correction approach, which leverages positional information from individual measurements and corresponding imaging data to estimate gene expression levels. By incorporating external latent informative cues, the Sprod correction method effectively enhances SRT noise reduction. Another notable approach is the SpotClean^7^ model, which models the diffusion pattern in SRT data using a Poisson distribution. It quantifies the extent of spot swapping between tissue spots and background spots to adjust unique molecular identifier (UMI) counts. This method allows for the accurate adjustment of UMI counts by accounting for the phenomenon of transcript swapping, thereby improving the quality of SRT data. Overall, these methods specifically tailored to address the unique challenges of SRT data have shown promising results in reducing noise and enhancing the accuracy of spatial transcriptomics analyses^9^.

The fundamental concept behind these recent methods involves fitting the diffusion patterns observed in SRT data through statistical distribution models and adjusting UMI counts to obtain denoised SRT data, thus enhancing downstream analysis. However, the phenomenon of transcript spot swapping is a random and directionless process heavily influenced by experimental manipulations, making it challenging to mathematically model the diffusion pattern accurately. Although applying force-fitting algorithms, as mentioned above, can lead to an increased false positive rate in SRT data, it is disheartening that the information from a subset of “valid genes” with low expression and spatial aggregation can be overshadowed. Furthermore, the evaluation methods employed for existing imputation techniques have predominantly focused on their capacity to recover true signals from noisy data, often neglecting the potential introduction of erroneous signals into the dataset.

Furthermore, the presence of experimental noise and missing data can obscure the underlying network structure, posing challenges to the efficacy of downstream network-based analyses, particularly in the context of community detection. Consequently, the preprocessing stage of denoising assumes great significance in enhancing the performance of community detection algorithms such as Louvain and Leiden when applied to SRT data14. Henceforth, we classify genes exhibiting substantial spatial aggregation as the true signal within SRT data, while genes demonstrating high diffusion levels are identified as noise. By addressing the challenge of decreased SNR resulting from transcript diffusion in SRT data, we can enhance the overall SNR by excluding genes with elevated diffusion levels. And we demonstrate that this denoising process holds paramount importance prior to undertaking any downstream analysis.

## Results

### Overview of GF for denoising SRT data

To effectively address the intricate spatial noise present in SRT data, we developed a denoising algorithm called the gene filter (GF) denoising algorithm. This algorithm is specifically designed to mitigate the impact of noise by evaluating the diffusion pattern of each gene using optimal transport techniques **(Fig. 1a)**. The GF algorithm filters out “invalid genes” that exhibit wide and uniform expression across tissue regions but lack meaningful differential expression necessary for clustering, cell type annotation, and the identification of DEGs in **Fig. 1b**. Notably, the GF algorithm ensures that the UMI counts of raw sequencing data remain unmodified. It only focuses on filtering noise-associated UMI counts without introducing additional error information. The GF algorithm is evaluated by its performance in enhancing the identification of DEGs, improving the specificity of marker genes, and elevating the clustering precision and the accuracy of cell type annotation **(Fig. 1c)**.

**Fig. 1.**
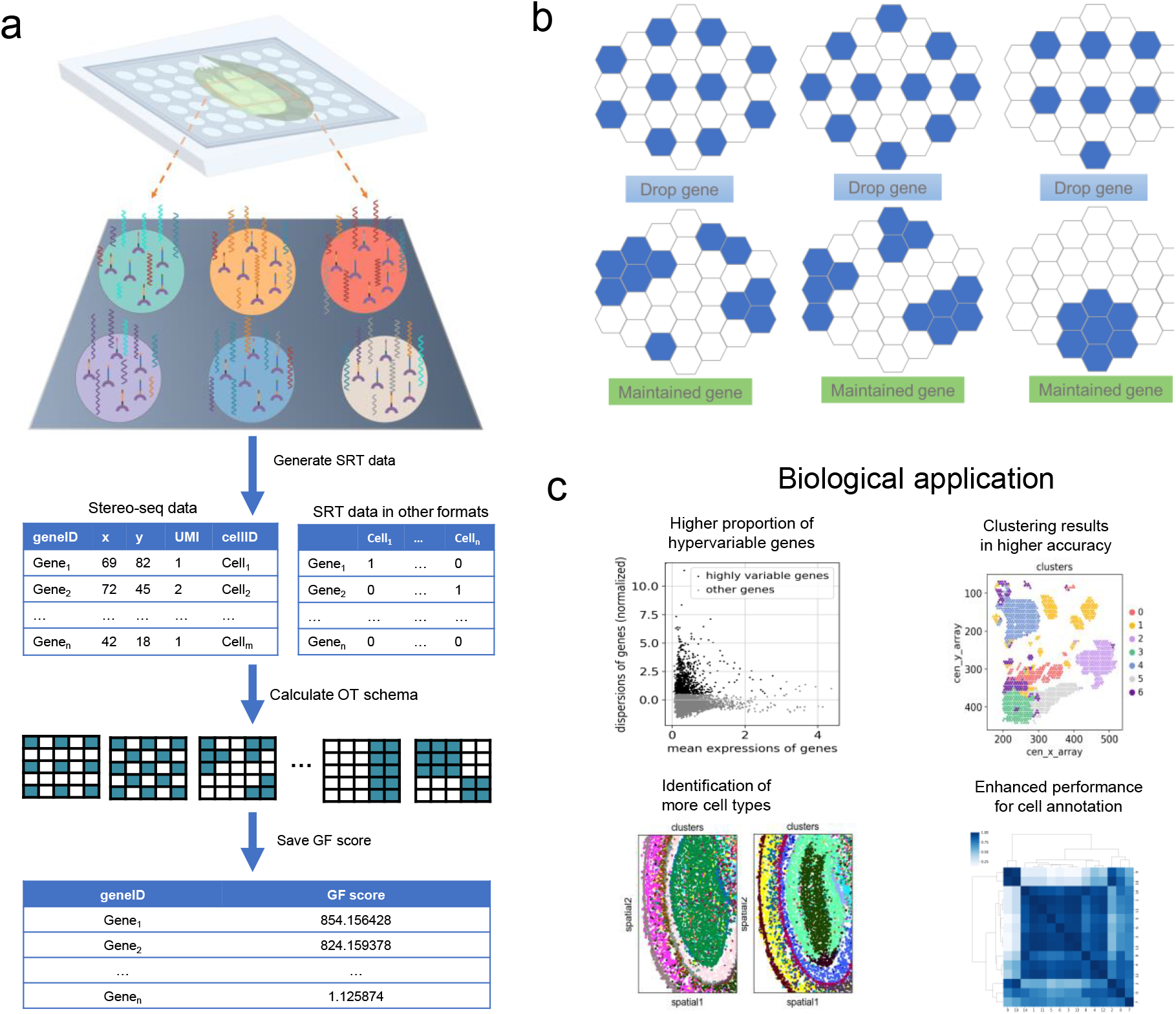
Overview of GF algorithm. **a**, The presence of transcript diffusion in SRT data is manifested by UMI counts in different formats of SRT data (10X Visium, Stereo-seq, etc.), and then the GF model is used to assess the degree of diffusion of each gene. **b**, Schematic diagram describing effective genes and the spatial distribution of effective genes. Blue honeycombs indicate UMI count present, whereas white honeycombs indicate UMI count absent. **c**, Four applications of denoised SRT data generated by GF for downstream analysis.

### SRT data contains noise that impedes downstream analyses

In our work, we focused on investigating noise phenomena in two prominent SRT technologies, namely 10X Visium and Stereo-seq^15^. Initially, we validated the diffusion of UMI counts in soybean root SRT data obtained using the Stereo-seq technology. Specifically, when utilizing bin 200 conditions (corresponding to cell-sized pseudo-spots measuring 100 μm square), we observed evident UMI counts extending beyond the actual tissue region, as illustrated in the FB staining image of soybean root tissue **(Fig. 2a)**.

**Fig. 2.**
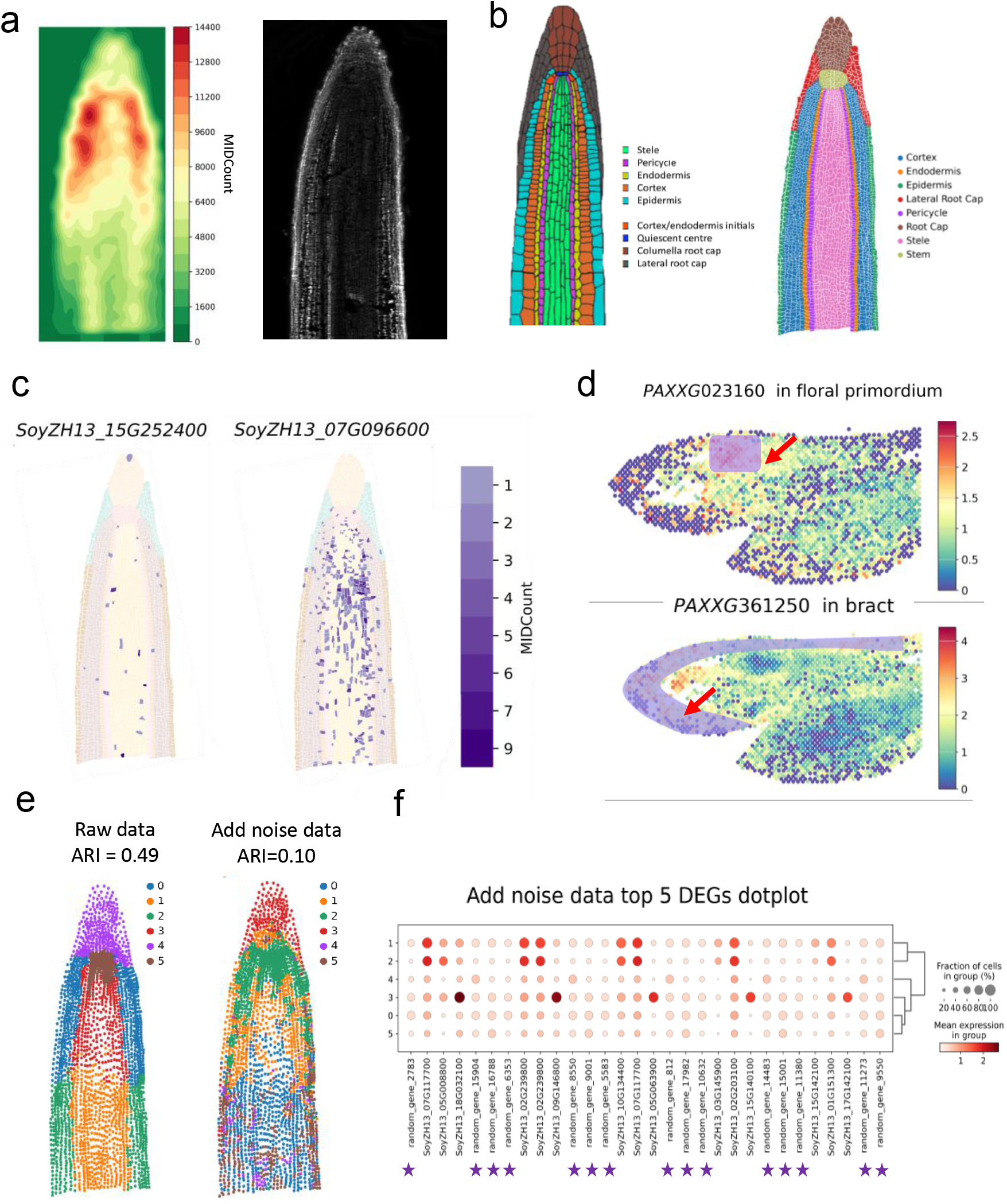
Noise in SRT data can interfere with subsequent analyses. **a**, Gene expression heatmap (bin size 200) of Stereo-seq soybean root longitudinal section data (left) and FB staining of tissue (right). **b**, Arabidopsis root cell type model diagram (left), and the main 8 cell types in the soybean root tissue (right) was produced as Ground truth. **c**, Gene expression heatmap of two marker genes *SoyZH13_07G096600* and *SoyZH13_15G252400* in the vascular bundle tissue in the actual Stereo-seq data. **d**, Gene expression heatmap of the marker genes *PAXXG023160* in floral primordium tissue and marker genes *PAXXG361250* in bract tissue, in the 10X Visium orchid slice data. **e**, Soybean root SRT data (left). Clustering results of raw SRT data (left) and SRT data (right) after adding random noise genes. **f**, Bubble chart of DEGs identified by clustering results of SRT data (right) after adding random noise genes. Only displayed the top 5 DEGs in each cluster.

Subsequently, we delved into the diffusion phenomenon within the soybean root tissue. We manually annotated eight major tissue regions (cortex, endodermis, epidermis, lateral root cap, pericycle, root cap, stele, and stem), enabling the creation of cell-bin (cell-sized pseudo-spots based on cell segmentation results) SRT data that closely resembled the actual situation **(Fig. 2b)**. Our analysis revealed that two marker genes, *SoyZH13_07G096600* and *SoyZH13_15G252400*^16^, which were expected to exhibit specific expression in vascular bundles, displayed broad expression throughout the entire soybean root tissue in the raw SRT data **in Fig. 2c**. This suggested that these marker genes undergone diffusion, leading to a decrease in their SNR and making it a challenge to characterize vascular cell types accurately. In addition, we also observed spatial drift in the expression patterns of marker genes across three slides of 10X Visium orchid flowers data. For instance, marker genes *PAXXG023160* and *PAXXG009220*^17^, which were expected to show specific expression in the primordium tissue **(Fig. 2d and Extended Data Fig. 1a)**, exhibited widespread and extensive expression outside the primordium tissue. Similarly, marker genes *PAXXG361250* and *PAXXG024270*^17^, associated with the bract tissue, displayed significant UMI counts in the surrounding areas of the bract tissue and even outside the tissue **(Fig. 2d and Extended Data Fig. 1a)**. This phenomenon was consistently observed in the other two sets of 10X Visium SRT data from orchid slides (slide 2 and slide 3) (**Extended Data Fig. 1b)**. Moreover, we observed a similar *exsitu* expression capture phenomenon in animal SRT data, such as the 10X Visium human prefrontal cortex SRT data (LIBD_151507)^18^. Differential marker genes for white matter (WM) tissue, including *GFAP* and *MOBP*, exhibited widespread expression outside the WM tissue (**Extended Data Fig. 1b)**. Collectively, these findings provide compelling evidence that the diffusion of transcripts introduces significant noise into SRT data.

Next, we confirmed that the noise in SRT data interferes with downstream clustering and the identification of DEGs. Utilizing the soybean root data, which had been annotated with cell type information, we applied SpaGCN software^19^ to cluster the raw SRT data consisting of 27,878 genes into six clusters. The clustering results achieved an Adjusted Rand Index (ARI)^20, 21^ value of 0.49, Accuracy (ACC) value of 0.66, Normalized Mutual Information (NMI)^21, 22^ value of 0.54, and Fowlkes-Mallows Index (FMI)^21^ value of 0.66 in **Fig. 2e**. To assess the effect of noise, we introduced 5,000 random noise genes under the same processing conditions (**Extended Data Fig. 1c)**. The clustering results with the added noise genes exhibited an ARI value of 0.10, ACC value of 0.47, NMI value of 0.18, and FMI value of 0.43 in **Fig. 2e (Supplementary table1, Supplementary table2 and Supplementary table3)**. These findings clearly demonstrate that the presence of widely expressed genes in the tissue significantly impairs the performance of clustering analysis. Moreover, we discovered that these noise genes exhibit strong specificity when identifying DEGs in **Fig. 2f**, highlighting the ineffectiveness of traditional differential expression analysis algorithms in effectively excluding such noise genes. Hence, performing effective denoising of SRT data becomes necessary prior to conducting downstream bioinformatics analysis.

### SRT data denoising algorithm GF based on optimal transmission

In this work, we present the GF algorithm as a novel approach for denoising SRT data. Different from conventional denoising algorithms that rely on mathematical statistical models to do imputation, GF focuses on gene filtering to remove true negative signals from the SRT data while preserving the heat map of the raw sequencing data. By effectively eliminating spatial noise signals and emphasizing “valid genes” with low diffusion levels, GF significantly enhances downstream bioinformatics analysis. Firstly, the GF leverages the positional information and UMI counts of spots in the SRT data to create a two-dimensional spatial distribution for each captured gene **(Fig. 3a)**. To estimate the outer contour of the tissue, we employ the alpha-shape algorithm^23^. Based on the source distribution characteristics and the proximity to the contour, a consistent target distribution is generated for each gene, representing its state of widespread expression throughout the tissue under maximum diffusion conditions. Subsequently, the optimal transportation method^24, 25^ is employed to compute the Kantorovich relaxation between the source and target distributions in **Fig. 3a**. During the computation of the minimum transportation cost, the GF sets a maximum iteration limit of 300,000 iterations to ensure reliable convergence (for detailed steps, please refer to the methods section). Following the same steps, GF calculates the optimal solution for each gene, representing the minimum transportation cost between the gene’s source distribution and its corresponding target distribution. This solution accurately reflects the diffusion extent of gene expression within the tissue, where a smaller GF value indicates a larger diffusion extent and vice versa. To generate denoised SRT data, a filtering threshold is determined based on the distribution of divergence coefficients for all genes, taking into account the overall diffusion extent of the SRT data. An automated algorithm based on gradient changes is provided to determine the filtering threshold, ensuring convenience and accuracy. Additionally, researchers have the option to manually set a filtering threshold that aligns with the characteristics of their specific SRT data. Furthermore, genes with global expression patterns within the tissue, such as housekeeping genes, are also assessed as high diffusion genes. To prevent their interference with subsequent clustering and differential gene expression calculations, GF typically filters out these genes. By removing such genes from the analysis, GF focuses on capturing genes with more localized and informative expression patterns, which can enhance the accuracy of downstream analyses.

**Fig. 3.**
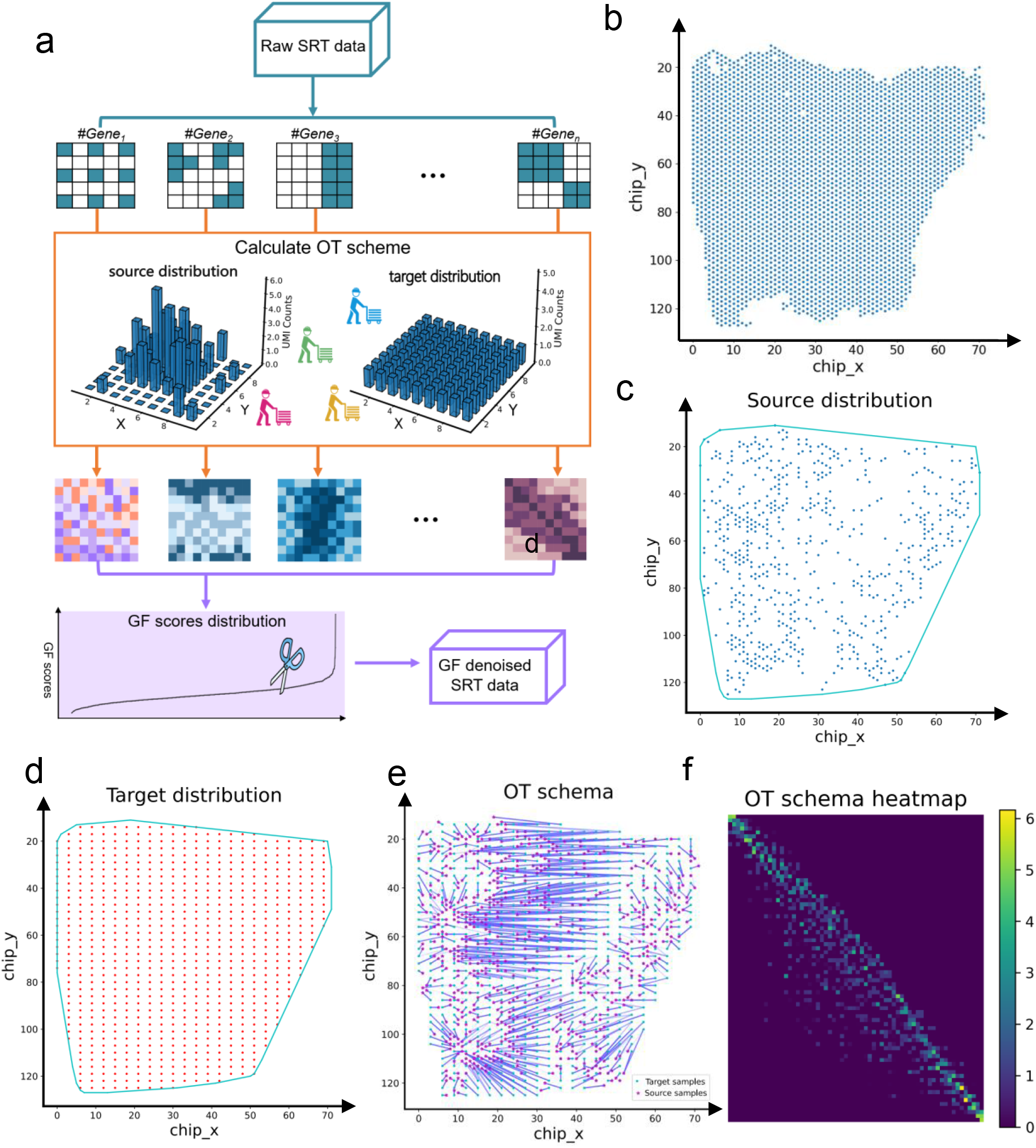
GF algorithm principle. **a**, Cartoon describing of GF model, from data preparation, feature extraction and calculate GF scores to SRT data denoising. **b**, All cell in 10X Visium colorectal cancer, and the x-axis and y-axis represent the spatial position of each cell. **c**, Expression model of gene *AC004556*.*3* in SRT data. The blue dots indicate the cell locations where the gene is expressed. The outline closed in cyan is the tissue boundary estimated using the Alpha-shape algorithm. **d**, The target distribution corresponding to the gene *AC004556*.*3* is generated by GF algorithm, and the sum of the source distribution and the target distribution UMI counts is equal. **e**, Optimal transfer scheme of Solution of Kantorovich Relaxation from the source distribution to the target distribution of gene *AC004556*.*3*. **f**, The heat map cost matrix of the source distribution and the target distribution of genes *AC004556*.*3*, and the optimal transmission scheme.

Moreover, we applied the GF algorithm to actual Stereo-seq data of soybean roots. As an example, we focused on the gene *AC004556*.*3* and analyzed its diffusion extent using the GF algorithm in human colorectal cancer SRT data **(Fig. 3b)**. First, we constructed the gene’s source distribution and target distribution **(Fig. 3c)**. Subsequently, we calculated the optimal transport solution, which provides insights into the gene’s diffusion extent, and visualized the optimal transport scheme from the source distribution to the target distribution **(Fig. 3d)**. In **Fig. 4e**, the red dots represent the 2D positions of all points in the source distribution, the cyan dots represent the 2D positions of all points in the target distribution, and the blue lines of different shades represent the cost of dot transport UMI counts from the dots in the source distribution to the target distribution. Additionally, **Fig. 4f** displays the matrix of the optimal transmission scheme. Rows in the matrix correspond to the source points of the source distribution, columns represent the target points of the target distribution, and each element value represents the quality of the transfer volume from the corresponding source point to the target point. For a more comprehensive understanding of how GF calculates the optimal solution, please refer to the methods section for detailed information.

**Fig. 4.**
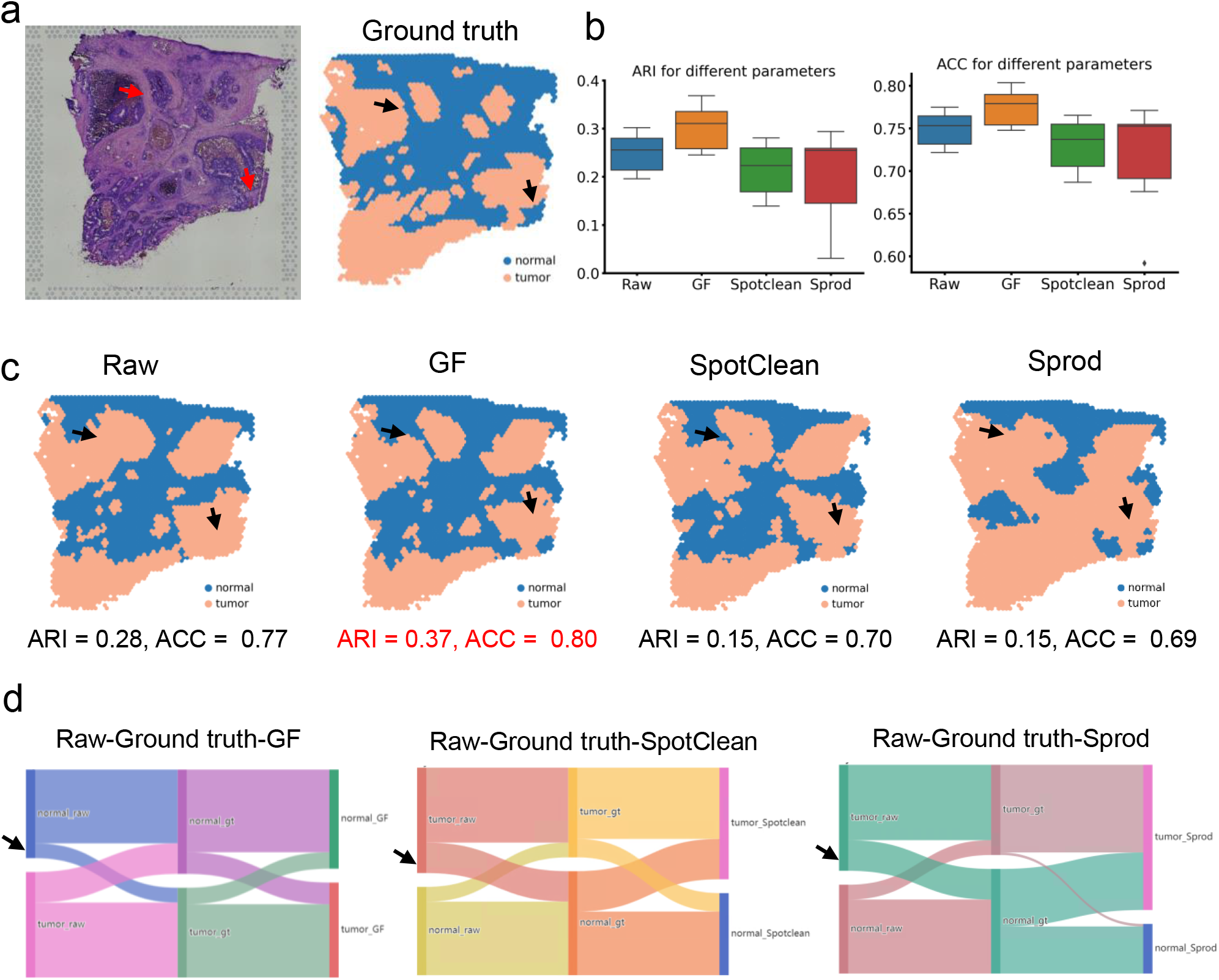
Identification of tumor cell types in 10X Visium colorectal cancer data is more accurate with GF-denoised SRT data. **a**, 10X Visium colorectal cancer section H&E staining map (left) and the distribution of tumor cells and normal cells from pathologist’s visual inspection (right). Orange dots represent tumor cells and blue dots represent normal cells. **b**, Distribution of ARI value and ACC value of raw, GF-denoised, SpotClean-denoised, and Sprod-denoised SRT data clustering results under 9 different clustering parameter conditions. Box boundaries represent interquartile ranges, whiskers extend to the most extreme data point (no more than 1.5 times the interquartile range) and the line in the middle of the box represents the median. **c**, Clustering results of Raw, GF-denoised, SpotClean-denoised, and Sprod-denoised SRT data, using SpaGCN software, set the number of clusters to 2, and the learning rate to 0.01. **d**, Three Sankey diagrams are the data flow between GF-denoised and raw and Ground truth, between SpotClean-denoised and raw and Ground truth, and between Sprod-denoised SRT data and raw and Ground truth.

In addition, to validate the accuracy of the divergence coefficient computed by GF in representing the diffusion level of gene two-dimensional distributions, we generated three sets of two-dimensional spatial Gaussian distributions with varying diffusion levels. Each set consisted of 10,000 spots, with spot numbers of 500, 1000, and 2000, respectively (**Extended Data Fig. 2, Supplementary table4 and Supplementary table5)**. Subsequently, we calculated the GF results for these distributions and observed a strong one-way correlation between the diffusion characteristics of the two-dimensional spatial distributions and the computed GF results, once random interference factors were excluded from the Gaussian distributions **(Extended Data Fig. 3a)**. However, the Moran’s I value^26^, which evaluates gene spatial autocorrelation, was found to be ineffective in characterizing the diffusion properties of these distributions **(Extended Data Fig. 3b)**. Furthermore, to exclude the influence of spot count which varies greatly for each gene, we generated 10,000 diffusion patterns with identical characteristics but with different numbers of spots (ranging from 2 to 10,002 with a step size of 1). In the case of two-dimensional spatial distributions with more than 10 points, the GF results exhibited periodic fluctuations while consistently remaining within the range of 3.7-3.8 **(Extended Data Fig. 3c, Supplementary table5)**. Although we expected the GF results to remain constant, we consider a fluctuation of 0.1 to be acceptable due to the inherent random nature of the Gaussian distribution generation process.

In summary, the experimental results demonstrate the ability of the GF algorithm to accurately assess the diffusion degree of genes across different numbers of expressed spots. This aligns with our objective of preserving genes expressed in fewer cells and with lower UMI counts.

### GF improves the accuracy of human colorectal cancer tumor cell identification

Pathologists commonly rely on H&E images to evaluate tumor extent and invasiveness. SRT data presents a distinctive opportunity to integrate pathological images with transcriptomic expression information, thereby enhancing tumor diagnosis and enabling precision treatment approaches.

First, we obtained the Ground truth SRT data **(Fig. 4a)** by integrating H&E staining images with the publicly available 10X Visium human colorectal cancer dataset^7^, which includes annotations of tumor and normal cell types for each spot. To enhance the discrimination between tumor cells and normal cells in a binary classification task, we utilized the SpaGCN software and explored nine different parameter settings. For each parameter set, we computed the clustering results of the raw, GF, SpotClean, and Sprod groups of SRT data. The selected parameter configurations included learning rates of 0.1, 0.01, and 0.001, and fixed cluster numbers of 2, 3, and 4**(Supplementary table4 and Supplementary table6)**. These settings were carefully chosen to strike a balance between achieving high prediction accuracy and preventing overfitting. Test results demonstrated that the GF-denoised SRT data yielded the best performance, as indicated by the distribution of ARI, ACC **(Fig. 4b)**, NMI, and FMI values **(Extended Data Fig. 4a)**, which serve as reliable evaluation metrics for assessing clustering results. In addition, we performed clustering validation on the same dataset using identical parameter settings. Among the various data types, including raw data (ARI=0.28, ACC=0.77), GF-denoised SRT data (ARI=0.37, ACC=0.80), SpotClean-denoised SRT data (ARI=0.15, ACC=0.70), and Sprod-denoised SRT data (ARI=0.14, ACC=0.69), GF demonstrated the highest accuracy in accurately identifying tumor cells in **Fig. 4c and Supplementary table7**. Furthermore, the Sankey diagram and confusion matrix revealed that GF and the Ground truth exhibited the least data flow error for tumor cells in **Fig. 4d and Extended Data Fig. 4**.

Furthermore, we generated heatmaps depicting the distribution of total expression level per cell and the distribution of the number of genes expressed per cell for the raw, GF-denoised, SpotClean-denoised, and Sprod-denoised SRT data **(Extended Data Fig. 5a)**. When observing the heatmaps with the color bar range set from the maximum to the minimum values, several observations can be made. Firstly, the heatmap of the total expression level distribution per cell shows that the GF-denoised SRT data exhibits the most similar distribution to the raw SRT data, while the Sprod and SpotClean denoising methods alter the expression distribution characteristics of the raw SRT data. Furthermore, although the GF denoising algorithm reduces the total expression per cell after filtering out “invalid genes,” the distribution characteristics of the expression heatmap remain consistent with the raw SRT data. This indicates that the “invalid genes” removed by GF do not exhibit significant spatial aggregation and are likely to represent genes with widespread expression across the tissue. Additionally, when examining the heatmap of the distribution of the number of genes expressed per cell, it becomes apparent that compared to the raw SRT data, the GF-denoised SRT data shows an increased proportion of cells with a higher number of genes, indicated by the intensified red-colored region. This provides further evidence that GF filtering specifically targets genes that are highly dispersed. In contrast, the results of the Sprod and SpotClean methods do not exhibit this characteristic in **Extended Data Fig. 5b**.

**Fig. 5.**
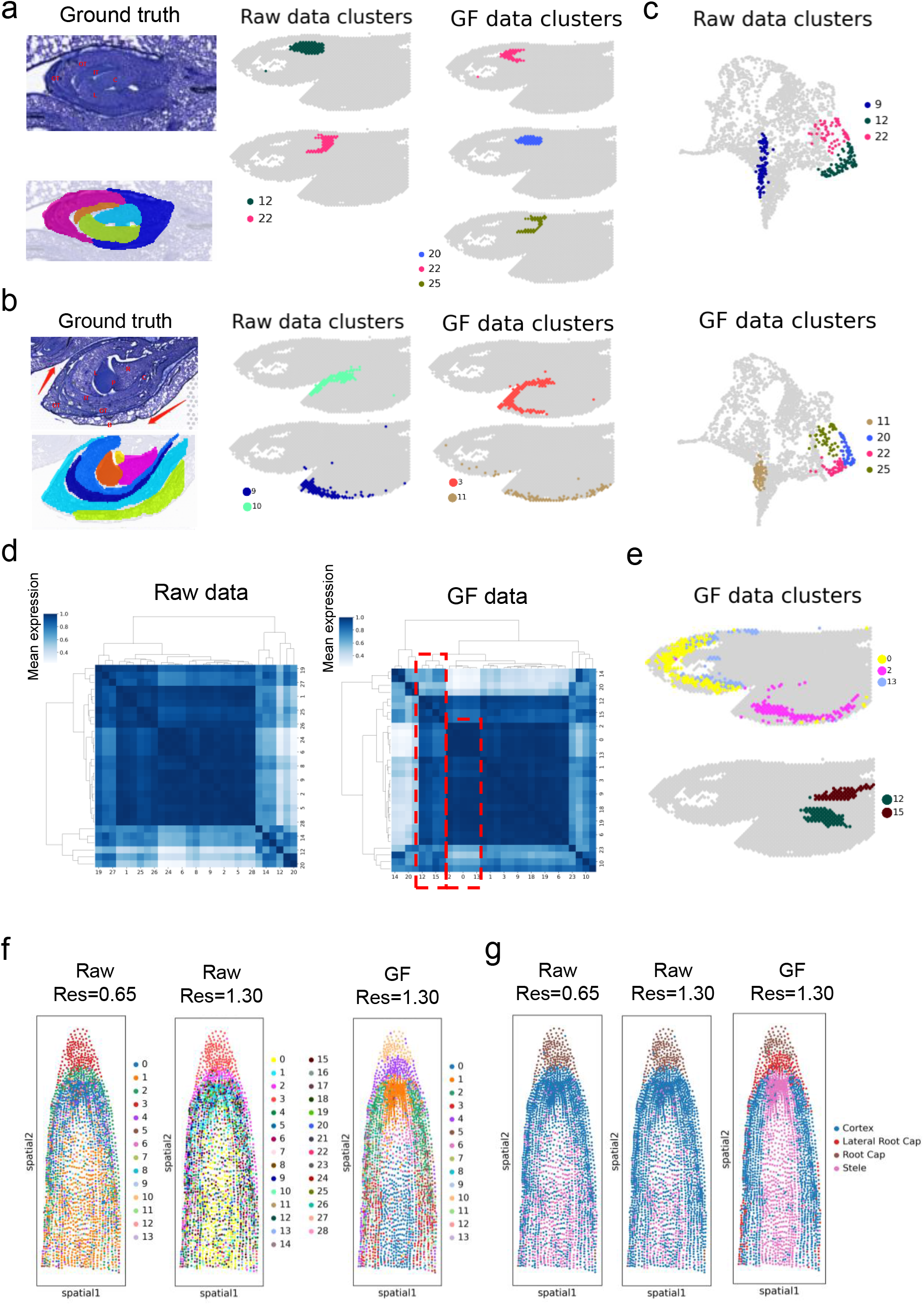
GF assists 10X Visium orchid dataset to identify more cell subtypes. **a**, In orchid bud region 1, two cell types were identified in the raw SRT data (clusters 12 and 22) and three cell types were identified in the GF-denoised SRT data (clusters 22, 20, 25). The left panel is the HE staining picture of bud 1 and the corresponding part of the Ground truth cell types. **b**, In orchid bud region 2, two cell types (clusters 10 and 9) were identified in the raw SRT data and two cell types (clusters 3 and 11) were identified in the GF denoised SRT data. The left panel in a and b is the H&E staining image and the cell types pattern picture corresponding to the flower bud. **c**, UMAP plot in clustering results of raw SRT data and GF-denoised SRT data. **d**, Cluster-cluster correlation heatmap of raw SRT data, and GF-denoised SRT data. The blue color bar indicates the UMI counts. **e**, In the clustering results of the GF-denoised SRT data, the two similar clusters 12 and 15 are observed together and found to be Bract tissues, and the three similar clusters 2, 0, and 13 are observed together and found to be near the end of the organization. **f**, Clustering results of soybean Stereo-seq data in Fig1 using Scanpy software. The clustering results of the raw SRT data and the GF-denoised SRT data when setting the same parameters (resolution=1.3, pc=30). When the raw SRT data set resolution=0.65, the clustering results of 14 clusters were obtained. **g**, Redistribute the three clustering results in Fig5. f according to the predicted cell type ratio in each cluster to form a new prediction result.

### Clustering results are more accurate with GF-denoised SRT data

Accurate cell type annotation serves as a critical initial step in unraveling the intricate process of early orchid bud development, encompassing inflorescence meristem, floral primordia, and organ identity determination. In this study, we assessed the enhanced clustering performance and the facilitation of identifying additional cell types by employing the GF algorithm on three publicly available 10X Visium datasets of orchid flowers^17^.

To determine optimal parameter settings in the absence of ground truth, we utilized the Scanpy software^27^ to explore a range of 60 parameter combinations. These combinations included resolutions spanning from 0.1 to 3.1 (with a step size of 0.1) and 30 or 50 principal components (pc) **(Supplementary table4 and Supplementary table6)**. Through this exploration, we observed that when employing a resolution of 3.0 and 30 principal components, the raw SRT data yielded 29 clusters, whereas the GF-denoised SRT data generated 30 clusters. This finding indicates that the GF algorithm is capable of identifying additional distinct cell subtypes, thus leading to improved clustering performance (**Extended Data Fig. 6a for visual representation**).

**Fig. 6.**
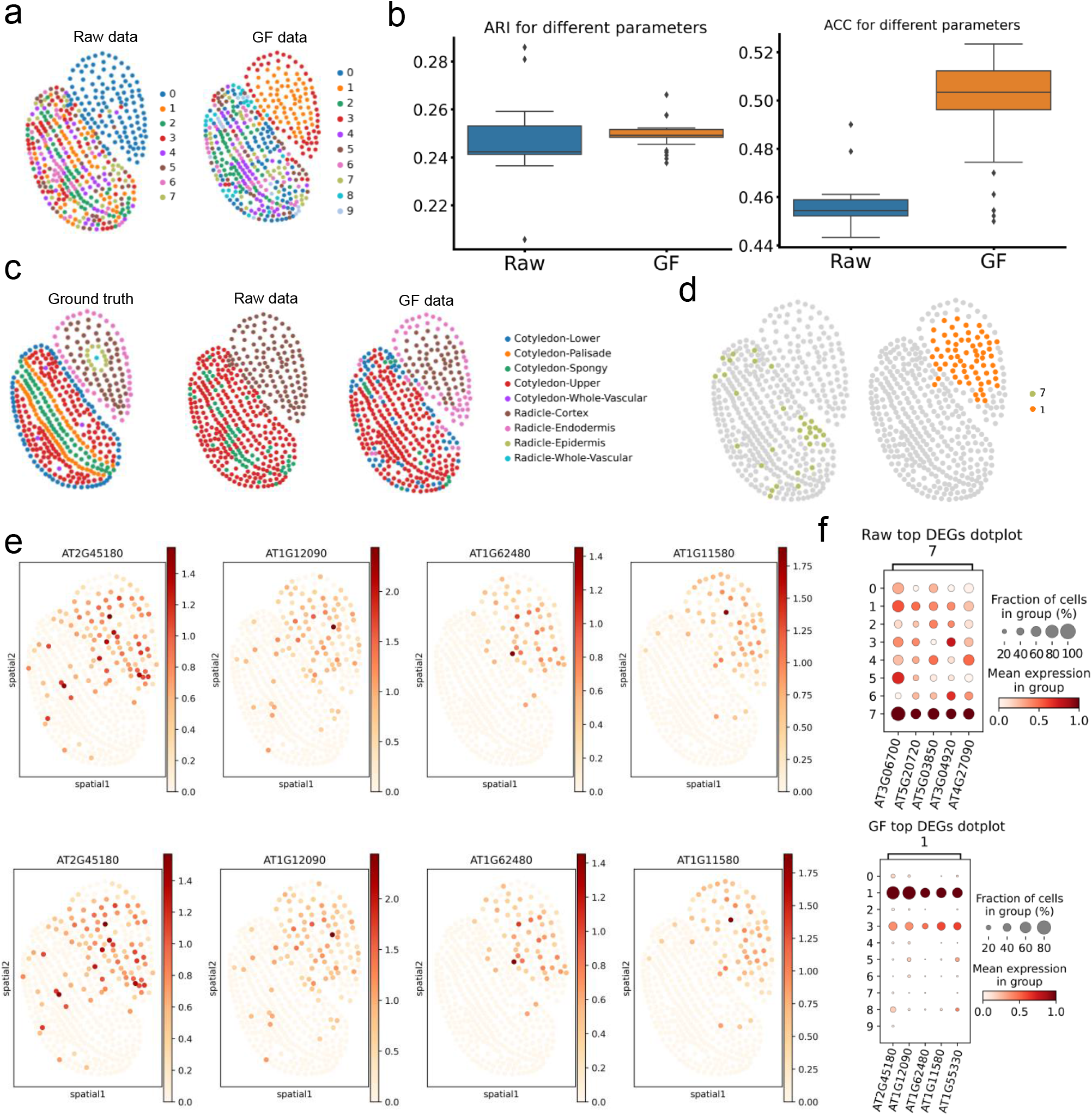
Identification of Stereo-seq Arabidopsis cotyledon and root tissue cell types is more accurate with GF-denoised SRT data. **a**, Clustering results of raw SRT data and GF-denoised SRT data. Set the number of clusters to 10 and the learning rate to 0.01 under the SpaGCN software clustering parameter conditions. **b**, The boxplot of the clustering results of ARI and ACC values for the raw SRT data and GF-denoised SRT data under 40 different parameter conditions using SpaGCN software. **c**, Left picture shows the Ground truth of Arabidopsis Stereo-seq data cell types. Calculated the contingency table for each cluster cell types and reassigned the categories, raw SRT data can only identify 3 cell types and GF-denoised SRT data can identify 5 cell types. **d**, Find the category with the highest clustering accuracy, the raw SRT data is cluster 7, the GF-denoised SRT data is cluster 1, and show the clustering results of these two clusters. **e**, Top four DEGs expression graph found by the cluster in Fig6. d, DEGs in cluster 7 is showed in the top four graphs, DEGs in cluster1 is the bottom four graphs. **f**, Bubble plots of the top 5 DEGs of cluster 7 and cluster 1 in Fig6. d.

In the clustering results depicted in **Extended Data Fig. 6a**, a detailed analysis of the floral bud region revealed notable differences between the raw SRT data and the GF-denoised SRT data. Specifically, when considering the tissue location information derived from the H&E staining image and the analysis of different cell type regions, the raw SRT data clustering only managed to identify the entire bud (cluster 12) and the meristematic tissue (cluster 22). In contrast, the clustering result of the GF-denoised SRT data successfully identified additional regions, including the tepal region (cluster 22), the column and lip regions (cluster 20), as well as the meristematic tissue (cluster 25) **(Fig. 5a)**. Moreover, when focusing on the patterned regions of cell types within the floral bud region, the clustering results of the raw SRT data solely identified the outer tepal (cluster 10) and bract (cluster 9) regions. However, it is worth noting that the bract region did not align well with the Ground truth. Conversely, the clustering results of the GF-denoised SRT data successfully captured the outer tepal tissue (cluster 3) and the bract tissue (cluster 11) with greater accuracy, exhibiting closer proximity to the Ground truth **(Fig. 5b)**. This observation is further supported by the UMAP plot, where the GF-denoised SRT data showcased stronger aggregation of cells within the bract tissue **(Fig. 5c)**. In addition, we analyzed the cluster-cluster correlation heatmaps derived from the two clustering results, which were calculated based on the expression differences between the two groups of clusters. The findings revealed that the heatmap generated from the GF clustering exhibited more discernible clustering patterns, making the GF results more advantageous for cell annotation in **Fig 5d**. Notably, clusters 0, 2, and 13 displayed close proximity to each other, indicating cell types within the tepal tissue, while clusters 2 and 15 were also closely associated, representing cells in the tail tissue **(Fig. 5e)**.

These observations further underscore the improved performance of GF in identifying distinct cell types and supporting more accurate cell annotation. We further assessed the enhanced clustering performance of GF on the slid 2 and slid 3 datasets of orchids. Clustering analysis was conducted using the Scanpy software for both datasets **(Supplementary table6)**. For slid 2, among the 60 sets of clustering parameters explored, the raw SRT data and the GF-denoised SRT data clustered into 20 clusters under the same parameter conditions (resolution=1.6, pc=30). However, the GF-denoised data exhibited a closer resemblance to the Ground truth, particularly in the clustering results of the outer tepal tissue **(Extended Data Fig. 6b)**. Moving on to slid 3, using the same parameter conditions (resolution=1.0, pc=50), the raw SRT data clustered into 14 clusters, whereas the GF-denoised SRT data yielded 12 clusters. However, the GF clustering results showcased improved performance in clustering the inner tepal and rostellum tissues. Notably, when analyzed with new parameters (resolution=1.5, pc=50), the GF-denoised SRT data also clustered into 14 clusters and also showed better performance than raw SRT data. In this scenario, the GF-denoised SRT data exhibited more accurate clustering outcomes for the inner tepal and Rostellum tissues in **Extended Data Fig. 6c and Extended Data Fig. 6d**.

Furthermore, we observed the clustering results are more accurate with GF-denoised SRT data in Stereo-seq soybean roots data. In the ground truth data of soybean roots, which consisted of eight cell types as shown in **Fig. 2b**, we utilized the Scanpy software to explore 78 different parameter combinations, including the resolution parameter range 0.1 to 4 with step 0.1, and pc of 30,50 **(Supplementary table8)**. The results clearly demonstrated that the clustering results obtained from the GF-denoised SRT data outperformed the raw SRT data across multiple evaluation metrics **(Fig. 5f, Extended Data Fig. 7a and Extended Data Fig. 7b)**. With the same parameter settings (resolution=1.30, pc=30), the clustering results for the raw SRT data clustered into 29 clusters and yielded ARI=0.10, ACC=0.48, NMI=0.19, and FMI=0.43, whereas the clustering results for the GF-denoised SRT data clustered into 14 clusters achieved ARI=0.40, ACC=0.65, NMI=0.40, and FMI=0.60. When analyzed with new parameters (resolution=0.65, pc=30), the raw SRT data also clustered into 14 clusters and achieved ARI=0.11, ACC=0.50, NMI=0.21, and FMI=0.47, showing worse performance than GF-denoised SRT data in **Fig 5f (Supplementary table9, Supplementary table10 and Supplementary table11)**. These findings clearly indicate that the application of GF denoising effectively enhanced the clustering performance of cell types **(Fig. 5f and Extended Data Fig. 7c)**.

Moreover, when reassigning categories based on the accuracy of predictions for each cluster, it became evident that the raw SRT data successfully identified only 3 distinct cell types, while the clustering results obtained from GF-denoised SRT data effectively distinguished 4 distinct cell types **(Fig. 5g)**. Overall, GF consistently demonstrated improvements in clustering performance across different parameter conditions.

### More accurate differential expression analysis with GF-denoised SRT data

Accurate identification of DEGs is crucial for unraveling key genes and cell fate determinants in Arabidopsis thaliana embryo root and leaf development research. The large field-of-view Stereo-seq technology enables the acquisition of subcellular resolution SRT data in Arabidopsis. By combining this data with spatial information, we are able to explore critical genes that play vital roles during the developmental processes of roots and leaves. Similarly, the Arabidopsis embryo Stereo-seq data **(Extended Data Fig. 8a)**, also exhibited the diffusion phenomenon between tissue and background in bin 30 SRT data.

First, we generated Arabidopsis thaliana Stereo-seq cell-bin data, with each sequencing unit representing a single cell. Collaborating with domain experts in biology, we annotated five cell types in the leaves and four cell types in the embryo roots based on the FB staining images **(Extended Data Fig. 8b)**, which served as the ground truth (Note: due to the small size of vascular cells, all vascular cells were merged into a single cell) **(Extended Data Fig. 8c and Supplementary table12)**. Subsequently, we utilized the SpaGCN software to perform clustering on both the raw SRT data and the GF-denoised SRT data, employing 41 different parameter combinations **(Supplementary table13 and Supplementary table14)**. The evaluation metrics, including ARI, ACC **(Fig. 6b)**, NMI, and FMI (**Extended Data Fig. 8c)**, consistently indicated the superior performance of the GF-denoised SRT data compared to the raw SRT data. Under the condition of setting the cluster number to 10 and the learning rate to 0.01, the raw SRT data could only cluster into 8 clusters, while the GF-denoised SRT data successfully clustered into 10 clusters. This is because the SpaGCN software merges similar clusters. Therefore, this improvement can be attributed to the GF algorithm effectively reducing noise in the SRT data, thereby unveiling previously concealed gene expression patterns in **Fig. 6b**.

In addition, by calculating the contingency table to determine the number of correctly predicted instances for each cluster compared to the Ground truth **(Supplementary Table 2)**, and reassigning categories based on the accuracy of predictions for each cluster **(Fig. 6c)**, we observed that the raw data identified only 3 different cell types, whereas the clustering results of the GF-denoised data successfully identified 5 different cell types. To further analyze the accuracy of predictions for each cluster in the clustering results **(Supplementary Table 2 and Supplementary Table 3)**, we selected the cluster with the highest accuracy. Cluster 7 exhibited the highest accuracy of 0.5625 for the raw data, whereas Cluster 1 achieved the highest accuracy of 0.7121 for the GF-denoised SRT data **(Fig. 6d and Supplementary Table 3)**.

We investigated the DEGs with strong specificity in cluster 7 and cluster 1, and their expression profiles were plotted **(Fig. 6e)**. The results revealed that the DEGs identified in cluster 7 of the raw data did not exhibit spatial expression patterns consistent with the cotyledon spongy tissue location. In contrast, the DEGs found in the radicle cortex tissue (cluster 1) of the GF-denoised SRT data displayed more accurate expression patterns, precisely reflecting the radicle cortex tissue region. Furthermore, the bubble plot of DEGs calculated from cluster 7 and cluster 1 demonstrates the increased significance of DEGs in the GF-denoised SRT data **(Fig. 6f)**. We also generated heatmaps illustrating the distribution of total expression in each cell and the distribution of the number of genes expressed per cell in the embryonic root and cotyledon **(Extended Data Fig. 8e)**. These heatmaps corroborate the findings from the 10X Visium colorectal cancer study, further supporting the effectiveness of the GF algorithm in removing noisy genes across different SRT data technologies.

Next, we compared the significance of DEGs identified by the t-test method between the clustering results of the raw SRT data and the GF-denoised SRT data in 10X Visium colorectal cancer data in **Fig. 4c**. For the reported 8 DEGs in tumor cells **(Extended Data Fig. 9a and Extended Data Fig. 10a)**, the GF-denoised SRT data exhibited larger fold change values and smaller P-value, indicating a higher level of significance in **Fig. 4c**. Similarly, for the 6 DEGs identified in normal cells, the GF-denoised SRT data showed larger fold change values and smaller P-value, indicating increased significance in **Fig. 4c (Extended Data Fig. 9b)**. Furthermore, we collected 6 tumor cell marker genes from the Cell Marker database and observed that their significance increased in the GF-denoised SRT data **(Extended Data Fig. 9c)**. Additionally, when considering the top 5 most significant DEGs calculated for tumor and normal cell types in each data group, the bubble plot clearly demonstrates that the GF results exhibit the highest similarity to the Ground truth **(Extended Data Fig. 10b)**.

In addtionally, we also observed the detection of DEGs is more accurate with GF-denoised SRT data in Stereo-seq soybean roots data. This improvement is further supported by the violin diagram of DEGs, where the DEGs identified in the GF results exhibited greater significance. Both clustering results accurately identified the root cap tissue, with the raw SRT data assigning it to cluster 3 and the GF-denoised SRT data assigning it to cluster 10. Therefore, we focused on the root cap tissue and compared the DEGs. It was found that the GF clustering results displayed larger fold changes and smaller p-value for the same set of DEGs **(Extended Data Fig. 9d)**, indicating stronger significance in the DEGs identified by GF. Furthermore, we generated violin diagrams illustrating the distribution of total gene expression per cell and the distribution of the number of genes expressed per cell for both the raw and GF-denoised soybean root data (**Extended Data Fig. 7a and Extended Data Fig. 7b**). The GF-denoised SRT data only reduces the overall expression level, but does not change the distribution of total count, indicating the “invalid genes” removed by GF.

These heatmaps provided additional evidence supporting the same conclusion observed in the Stereo-seq Arabidopsis root and leaf data, confirming the effectiveness of the GF algorithm in removing noisy genes in various plant SRT datasets.

## Discussion

Due to the necessary experimental procedures and the liquid experimental environment, part of the captured transcripts of individual genes in SRT data may not reflect their true *in situ* expression accurately. Consequently, the two-dimensional spatial distribution of gene expression loses its original spatial specificity, leading to a non-negligible decrease in SNR. Current solutions focus on adjusting the UMI counts within each spot to obtain interpolated or modified SRT data, aiming to enhance downstream analysis. Nevertheless, these methods encounter the limitation of fitting stochastic noise, which inherently lacks directionality with regular statistical models. Consequently, numerous false-positive data spots may be present in the SRT data, overshadowing the significant information on low-expression genes with spatial specificity.

To overcome this challenge, we developed the GF algorithm, which denoises SRT data effectively by eliminating the influence of true noise. The GF algorithm utilizes two-dimensional source distribution and target distribution for each gene. The source distribution represents the gene expression spatial distribution, while the target distribution is constructed as a uniform diffusion state which is the worst case. Different from conventional methods that compute the similarity between distributions, the GF algorithm assesses the distance between the two distributions accurately through an iterative process that computes the optimal solution in optimal transport.

The evaluation of the diffusion extent of individual genes enables identify “invalid” genes that cannot be highlighted by traditional SVG calculation methods (such as methods for calculating variance). Regarding the selection of the divergence coefficient filtering threshold, we have devised two approaches. The first approach leverages the gradient automatically generated from the distribution of divergence coefficients, providing a recommended threshold value for GF. This automated approach ensures ease of use for users. The second approach allows users to set their own filtering threshold, which can be specified as a percentage of genes to be filtered. This flexible option caters to the diverse preferences and requirements of users, accommodating SRT data with varying levels of diffusion. These two methods combine automation and user control, enabling efficient and adaptable processing of SRT data.

Furthermore, we conducted extensive validations to demonstrate the superior performance of the GF algorithm compared to other denoising algorithms specifically designed for SRT data, such as Sprod and SpotClean. We also verified that GF provides a more effective representation of gene diffusion compared to directly assessing the spatial autocorrelation (Moran’s I) of individual genes. One key advantage of the GF algorithm is its capability to accurately evaluate the diffusion coefficient of individual genes, even in the presence of variations in the number of expressed spots per gene. This feature enables GF to effectively filter genes exhibiting significant diffusion. Moreover, GF can be employed to enhance clustering algorithms by serving as a robust principal component or as a powerful tool for identifying spatially variable genes. Through these comprehensive validations, we have established that GF outperforms existing denoising methods, exhibits superior evaluation of gene diffusion, and offers versatility in improving downstream analyses. The GF algorithm represents a significant advancement in the field of SRT data analysis, providing researchers with a valuable tool for accurate gene expression profiling and spatial analysis. While our work has made significant advancements in characterizing gene diffusion using the GF algorithm, we acknowledge limitations in the automatic threshold recommendation algorithm. To address this, we utilized a method based on the gradient change in the one-dimensional distribution of GF results for all genes to select the filtering threshold. While this approach provides a reasonable starting point, it may not always yield the optimal denoising effect in every scenario. To offer greater flexibility and cater to specific requirements, we have designed an interface that empowers users to define their own filtering threshold. By allowing users to customize the threshold, they can fine-tune the denoising process and achieve the best results based on their unique experimental conditions and objectives. This user-defined threshold feature enhances the usability and adaptability of the GF algorithm, enabling researchers to optimize the denoising outcomes according to their specific needs.

The utilization of SRT data in addressing biological inquiries has gained significant momentum. However, the quality of SRT data poses a persistent challenge. Unlike the traditional drop-outs observed in scRNA-seq data, SRT data presents a more intricate spatial noise, further complicating the denoising process. The development of the GF denoising algorithm tailored specifically for SRT data represents a crucial initial stride in the processing pipeline for SRT data. By effectively addressing the inherent noise complexities of SRT data, the GF algorithm lays a solid foundation for downstream analyses and facilitates more accurate interpretations of the underlying biological phenomena.

## Methods

### Version

The following software and packages were used in the analysis: R-4.0.2; R/SpotClean0.99; R/Seurat-3.2.2; R/SPOTlight-0.1.7; Python-3.9.12; Python/Scanpy-1.9.2; Python/Anndata-0.8.0; Python/numpy-1.21.6; Python/pandas-1.3.5; Python/SpaGCN-1.2.5; Python/Louvain-0.7.1; Python/scipy-0.7.3; Python/UMAP; Python/Alphashape-1.3.1.

### Generation of soybean root and Arabidopsis SRT data

Initially, OCT-embedded blocks of soybean root tissues and Arabidopsis embryo were prepared for subsequent experiment. The tissues were precisely sectioned and underwent a series of essential procedures including fixation, staining, and imaging. To capture specific characteristics of the plant tissue sections, Calcofluor white staining image was employed to visualize the cell wall, while ssDNA staining image facilitated the visualization of nuclear patterns. Leveraging the CW-stained images, which corresponded to the sequencing data, we delineated the boundaries of individual cells and assigned cell type labels to each cell, thus creating the Ground truth SRT data. We utilized a combination of manual cell circling and automated cell segmentation software, specifically Cellpose, to accurately identify and delineate the complete cell wall curves for each individual cell. This approach allowed for precise characterization of the cellular boundaries in our analysis. The spatial position of each cell on the sequencing chip was represented by its centroid coordinates. With the assistance of the Ground Truth data, a comprehensive quantitative evaluation of the GF algorithm was conducted, allowing for a thorough assessment of its performance.

### GF denoising algorithm

GF denoising algorithm leverages optimal transmission theory to evaluate the diffusion degree of each gene of SRT data. To illustrate this, let’s consider a specific gene, gene P. The source distribution *P*(*x, y*) is constructed based on the gene’s transcript capture in the SRT data at 2D positions. It incorporates information from three dimensions: x, y, and UMI counts expression. Each coordinate point within the distribution represents the UMI counts for that gene. Next, a global uniform distribution *Q*(*x, y*) is constructed as the target distribution, representing the tissues with an equal number of points captured as gene P. The new spots in distribution Q are assigned the same UMI counts, ensuring that the sum of UMI counts in distribution Q matches the sum of UMI counts in gene P.

The transportation cost, denoted as *C*(*x*_0_, *y*_*o*_, *x*_1_, *y*_1_), represents the cost of moving 1 unit from a point *P*(*x*_0_, *y*_*o*_) in the source distribution P to a point (*x*_1_, *y*_1_) in the target distribution. A The cost function C is constructed using the squared Euclidean distance, defined as *C*(*x*_0_, *y*_*o*_, *x*_1_, *y*_1_) = (*x*_0_ − *x*_1_)^2^ + (*y*_0_ − *y*_1_)^2^. To establish the transport plan, T, we assign a weight, denoted as *T, T*(*x*_0_, *y*_*o*_, *x*_1_, *y*_1_) = *w*, w means the unit from (*x*_0_, *y*_*o*_) to (*x*_1_, *y*_1_). The transport plan T encompasses all possible starting positions and destinations. Currently, transport plan T must satisfy the following three conditions:

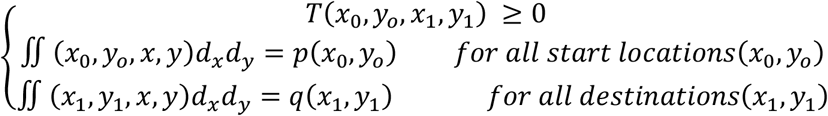

Hence, the total transportation cost from the source distribution *P*(*x, y*)to the target distribution *Q*(*x*_1_, *y*_1_) can be calculated as: 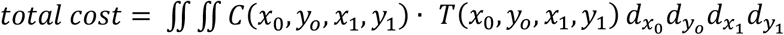. In the GF algorithm, the divergence coefficient of the gene under evaluation is transformed into a discrete optimal transport problem. or a gene P with n 2D positions capturing transcripts, the discrete positions can be represented as 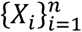, here, *δ*_*x*_ denotes the Dirac delta function *X ∈ R*^2^ at a 2D position, and the discrete distributions of source distribution and target distribution are 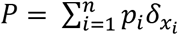 and 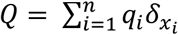. Enumerate the transportation cost of (*n*^2^ + *n*)*/* 2 for all Spot, *T ∈ R*^*n×m*^, then move one unit cost*C*_*ij*_ from the source distribution from *X*_*i*_ to *X*_*j*_ denoted as: *C*_*ij*_ = *‖X*_*i*_ – *X*_*j*_ ‖. Then the total cost of the transport plan can be expressed as: 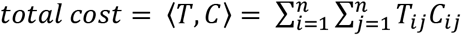.

By utilizing the Frobenius inner product between two matrices, the optimal transport plan can be reformulated as the following linear programming optimization problem:

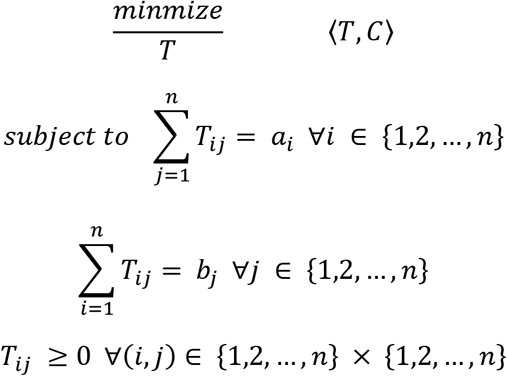

Denote *T*^∗^ as the solution of the optimization problem above, then the final calculated Wasserstein distance can be expressed as *W*(*P, Q*) = (⟨*T*^∗^, *C*⟩)^1*/*2^. The optimal transportation cost between the source distribution and target distribution of gene P can be calculated as *W*(*P, Q*). This distance reflects the proximity of gene P to the globally uniform distribution of tissues generated based on the P distribution in SRT data. A smaller *W*(*P, Q*) indicates a lower transportation cost, indicating a higher degree of diffusion for gene P. Conversely, a larger *W*(*P, Q*) signifies a higher transportation cost, suggesting a lower degree of diffusion for gene P. By evaluating the minimum transport cost *W*(*P, Q*) for all genes, noisy genes can be identified and screened.

In terms of time complexity, the computation speed of GF is relatively fast for technologies like 10X Visium, where the number of spots covering the entire tissue is typically not more than 10,000. However, for subcellular resolution spatial technologies, the number of spots capturing UMI counts for a single gene across the tissue can reach tens of thousands or more, resulting in slower computation speed for GF. To address this, we have introduced a solution based on the bin size parameter, which allows for the merging of adjacent spots into a single spot based on Euclidean distance. When adjusting the number of constructed source distribution points (500/1000/2000) while maintaining a constant degree of diffusion, the GF results consistently demonstrate a high level of consistency **(Extended Data Fig. 2a)**. This finding suggests that our down-sampling technique effectively reduces the number of spots.

### Target distributions in GF

In this work, we employed the Alpha-shape algorithm as a crucial step to accurately estimate the overall contour of the target tissue. The 2D Alpha-shape algorithm operates by rolling a circle with a radius of α around the given set of spots S in the SRT data. The points that intersect with the circle are identified as the edge contour points of S, while the trace of the rolling circle represents the boundary line or outer contour of S. This algorithm is particularly effective in capturing concave edges commonly found in tissue samples, providing an accurate estimation of the tissue’s outer contour. Importantly, since the outer contour can be directly derived from the SRT data itself, which inherently contains spatial information, the need for additional staining images is circumvented. This approach streamlines the process and eliminates the requirement for acquiring separate staining images.

For SRT data with a resolution that captures multiple spots per cell, such as 10X Visium data, the GF algorithm involves constructing both the source distribution and the target distribution for a specific gene. After generating the two-dimensional spatial distribution (source distribution), it is necessary to create a corresponding uniform distribution (target distribution) based on the number of expressing spots and the total UMI count information of that gene. Constructing the target distribution entails creating an evenly spaced distribution within the irregular tissue region while ensuring it contains the same number of spots as the source distribution. This approach guarantees that the UMI count values across all spots in the target distribution are equal and collectively sum up to the same total UMI count as the gene’s source distribution. By maintaining this balance between the source and target distributions, the GF algorithm effectively evaluates the diffusion degree of each gene in the SRT data.

For SRT data with subcellular resolution, where a single spot captures multiple cells, such as Stereo-seq data, the first preprocessing of SRT data involves incorporating tissue imaging information to retain only the expression information within the tissue region. In the case of cell-bin SRT data, when generating the source distribution for a specific gene, it involves calculating the center coordinates of each cell and the total sum of UMI counts within that cell. The corresponding target distribution for the gene is designed to have equal expression in each cell, ensuring that the total sum of UMI counts in the target distribution matches the total UMI count of that gene. The preprocessing steps for the raw SRT data closely resemble those of the 10X Visium SRT data. However, an additional step is introduced to scale the SRT data effectively. This involves defining a bin size, where each bin corresponds to a cell, and the center of each bin is assigned as the two-dimensional coordinates of the respective cell. By implementing this approach, we achieve a precise representation of gene expression at the subcellular level, enhancing the accuracy of subsequent analyses.

### Generation of random noise gene data

To assess the potential impact of “invalid” genes, which exhibit widespread and random expression in tissue regions, on downstream analysis, we conducted experiments using a dataset of soybean roots. This dataset consisted of 27,878 genes that were actually captured, along with 2,295 cells. To generate random noise genes, we followed a two-step procedure. Firstly, we randomly determined the number of cells in which the gene would be expressed by selecting a uniform distribution ranging from 20 to 2,295 cells using the Numpy library in Python. The centroids of these cells were recorded. Secondly, we assigned UMI count values to the gene in each cell by generating random integers from 1 to 5 using the “randint” function from the random library in Python. These integers represented the total UMI counts for the random gene in individual cells. This process was repeated to generate a total of 5,000 random noise genes. Subsequently, we compared the clustering results and differentially expressed genes between the original SRT data with 27,878 genes and the SRT data with added noise, resulting in 32,878 genes under the same experimental conditions. By conducting this comparison, we aimed to evaluate the impact of including these random noise genes on the analytical outcomes.

### Evaluation index of clustering results

This study employs four primary evaluation metrics to assess the clustering results: Adjusted Rand Index (ARI), Accuracy (ACC), Normalized Mutual Information (NMI), and Fowlkes-Mallows Index (FMI). ARI is utilized to measure the similarity between two clustering results, specifically comparing the clustering results of the raw SRT data and the ground truth, as well as the clustering results of the denoised SRT data and the ground truth. However, it is important to note that ARI values can be influenced by the number of cells present. ACC, on the other hand, quantifies the accuracy of the clustering result by comparing the spatial positions of cells in each cluster (designated as B) to the actual cell types in the corresponding clusters of the ground truth (designated as A). By recording the maximum count of correct cell type assignments (N1 to N6) in clusters C1 to C6, ACC provides a direct assessment of the consistency between the clustering result and the ground truth labels, enabling the calculation of the accuracy for each cluster prediction. NMI evaluates the degree of information shared between two clustering results, indicating their similarity, while FMI measures the similarity between clustering results based on the matching of data point pairs. These four metrics collectively offer a comprehensive and quantitative characterization of the clustering performance of the SRT data, providing valuable insights into its quality and accuracy.

### Generation of two-dimensional Gaussian distributions

To evaluate the ability of the GF algorithm to characterize the diffusion patterns of gene expression in a two-dimensional space, we utilized the estimated covariance matrix of a two-dimensional Gaussian distribution to assess the clustering degree of the distribution. Assuming the two-dimensional Gaussian probability density function as 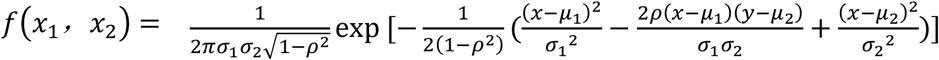, where 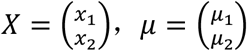 and the covariance matrix 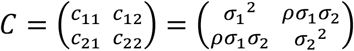, assuming 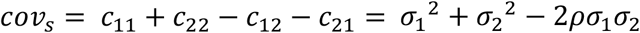 to represent the clustering degree of the two-dimensional Gaussian distribution. To evaluate the capabilities of the GF algorithm, we conducted experiments using two sets of simulated data. In the first set, we generated a series of Gaussian distributions with consistent diffusion degrees, ranging from 2 to 10002 points with a step size of 1. The mean was set as μ= [0,0], and the covariance matrix was defined as C=[[2,1], [1,2]]. Throughout these distributions, the *cov*_*s*_ value was fixed at 2. While the occurrence of points in these distributions introduced some level of randomness, the resulting fluctuations in diffusion patterns were deemed acceptable. This dataset was employed to validate the GF algorithm’s effectiveness in evaluating the diffusion degree of numerous genes with varying spot quantities. In the second set, we generated Gaussian distributions with diverse clustering degrees by altering the covariance matrix C as C=[[i,1], [1,i]], where i*∈*[1,10000]. We also assigned three different spot values (500, 1000, 2000) and computed the GF results for each dataset. This second dataset was specifically designed to demonstrate the GF algorithm’s ability to quantitatively assess the diffusion degree of individual genes based on their clustering characteristics. These two sets of simulated data served as valuable tools in evaluating and showcasing the GF algorithm’s performance in assessing diffusion patterns and quantifying gene expression at a spatial level.

## Supplementary information

Supplementary information includes 10 figures, and 14 tables.

## Acknowledgements

The author would like to thank reviewers for their reading and constructive comments.

## Author contributions

Lin Du and Bohan Zhang jointly developed the software. Lin Du contributed to algorithm verification, bioinformatics analysis, original manuscript writing, and figure generation. Jingmin Kang conducted the experiments and generated the Stereo-seq data. Haixi Sun and Bohan Zhang critically reviewed the manuscript and provided overall supervision for the study.

