## Supplementary material for "Optimal Transport Method-Based Gene Filter (GF) Denoising Algorithm for Enhancing Spatially Resolved Transcriptomics Data": Extended Data Fig.pdf

**Extended Data Fig.1**

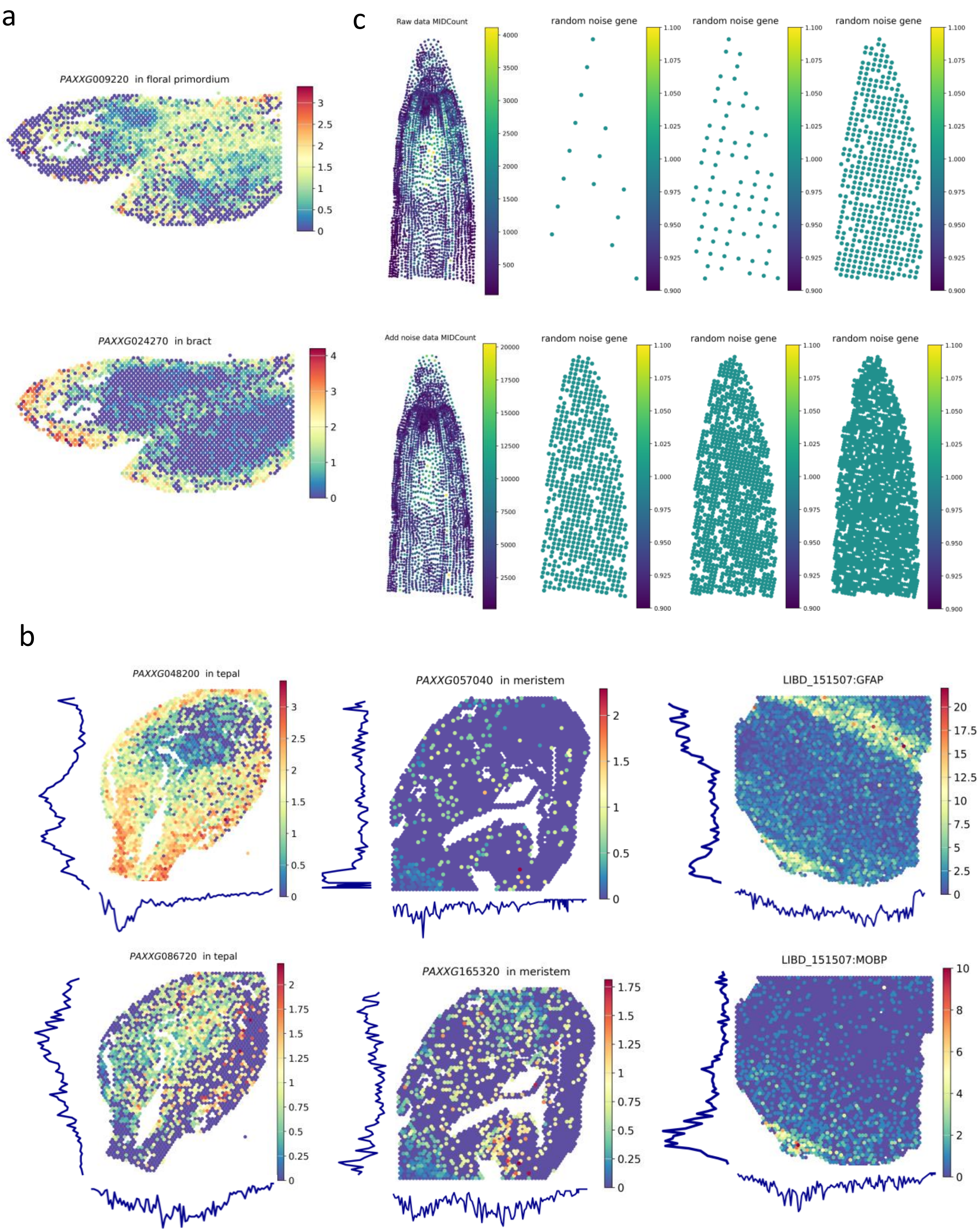

**Extended Data Fig.2**

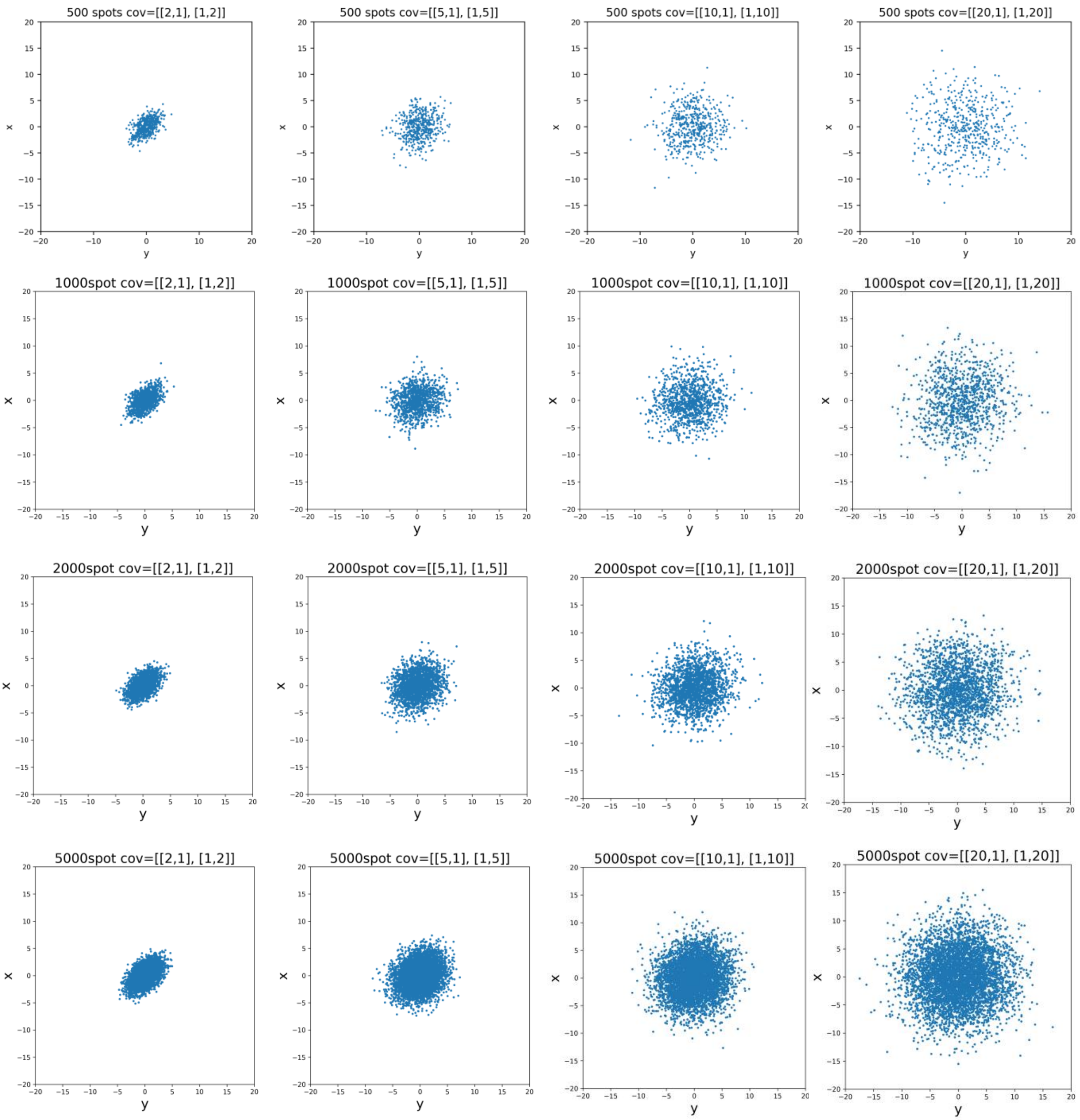

**Extended Data Fig.3**

a

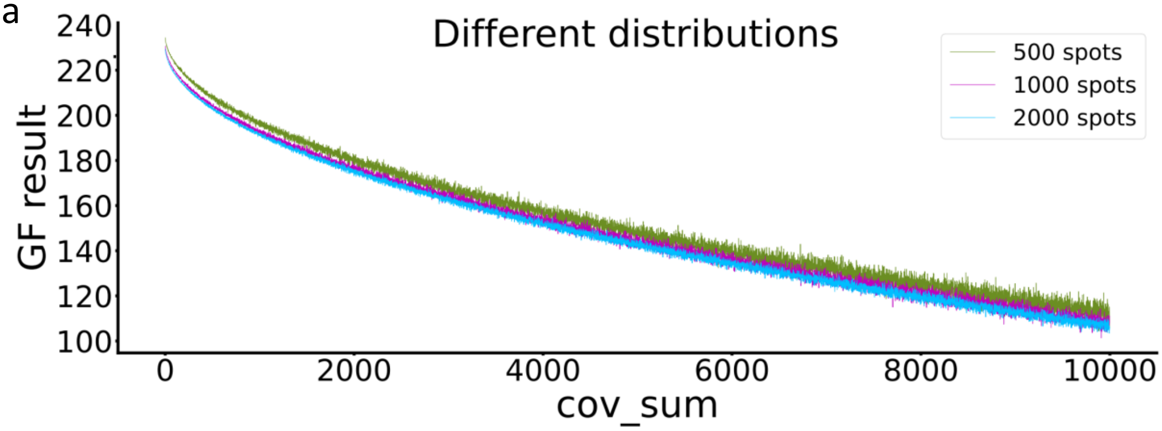


b

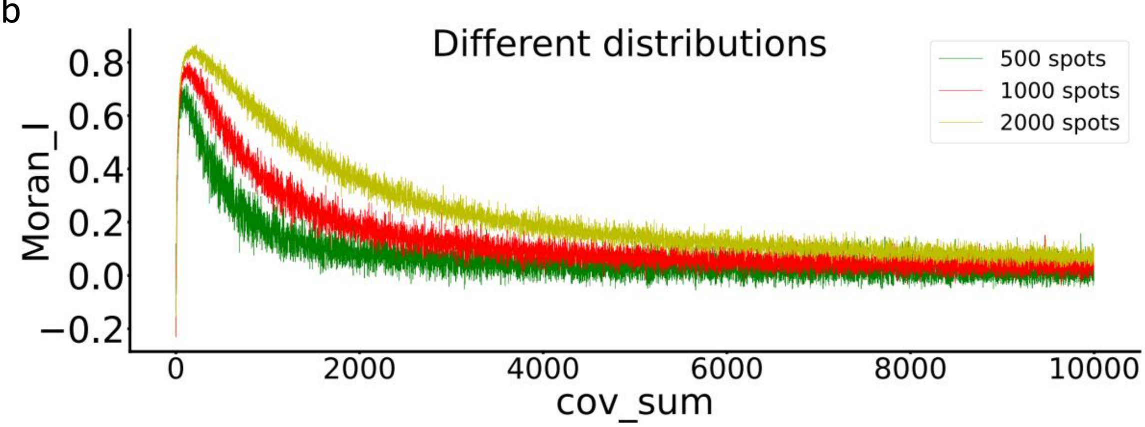


c

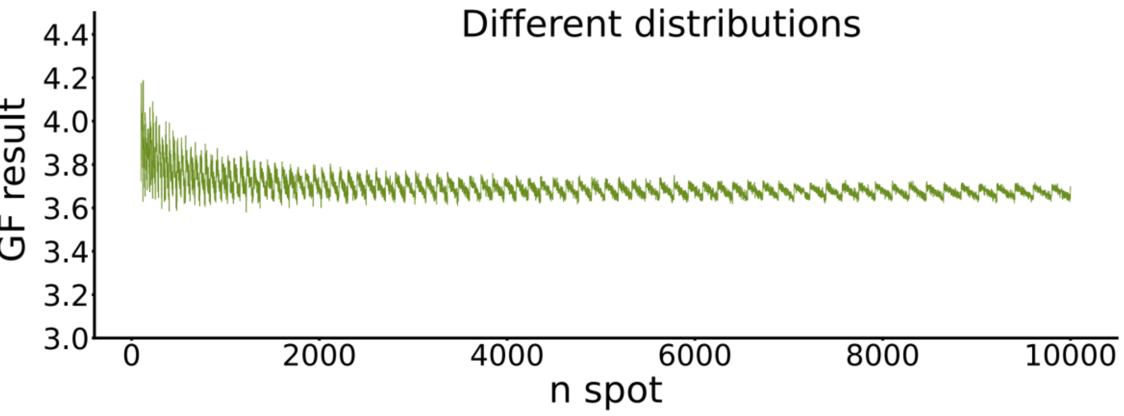

**Extended Data Fig.4**

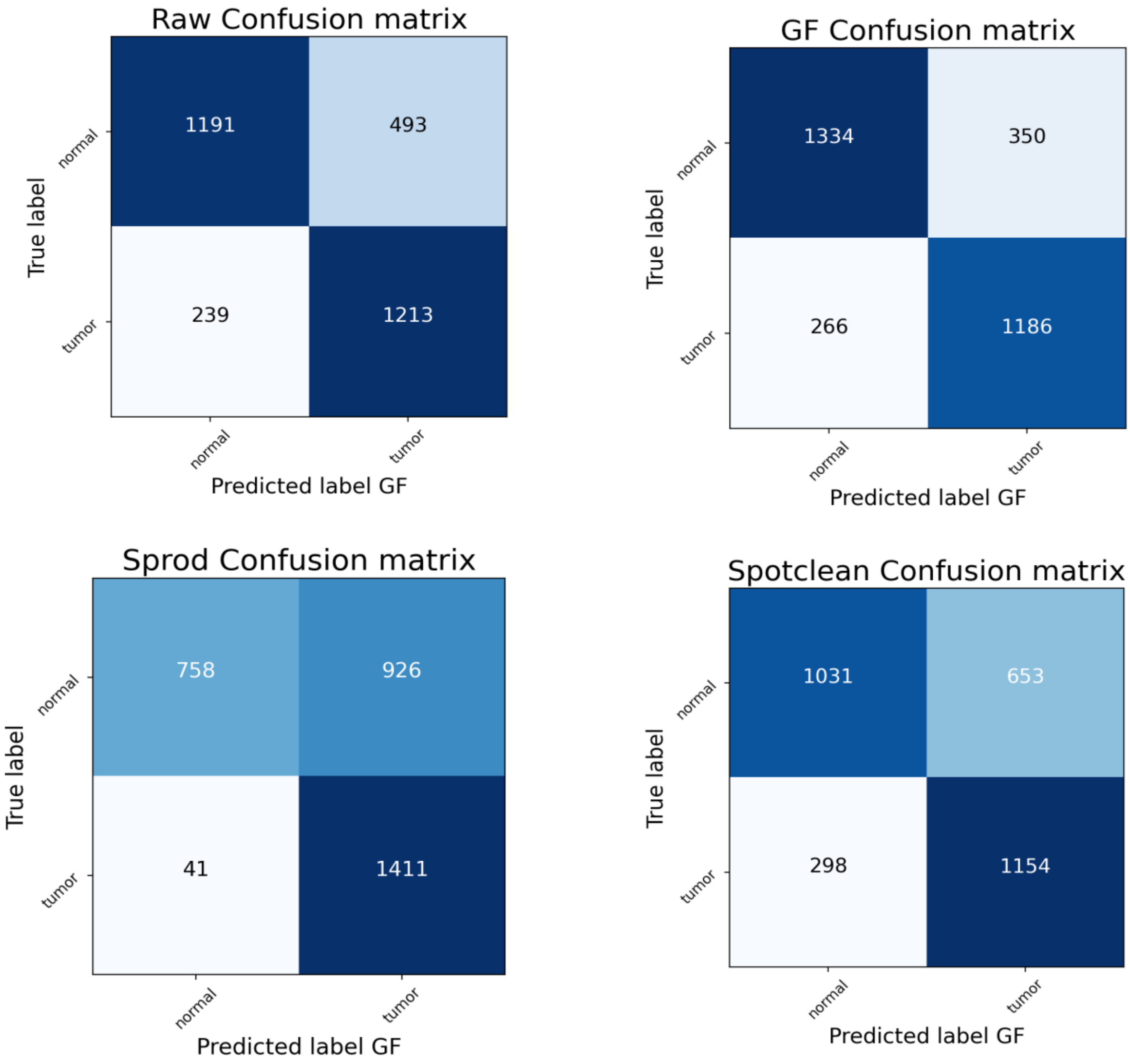

**Extended Data Fig.5**

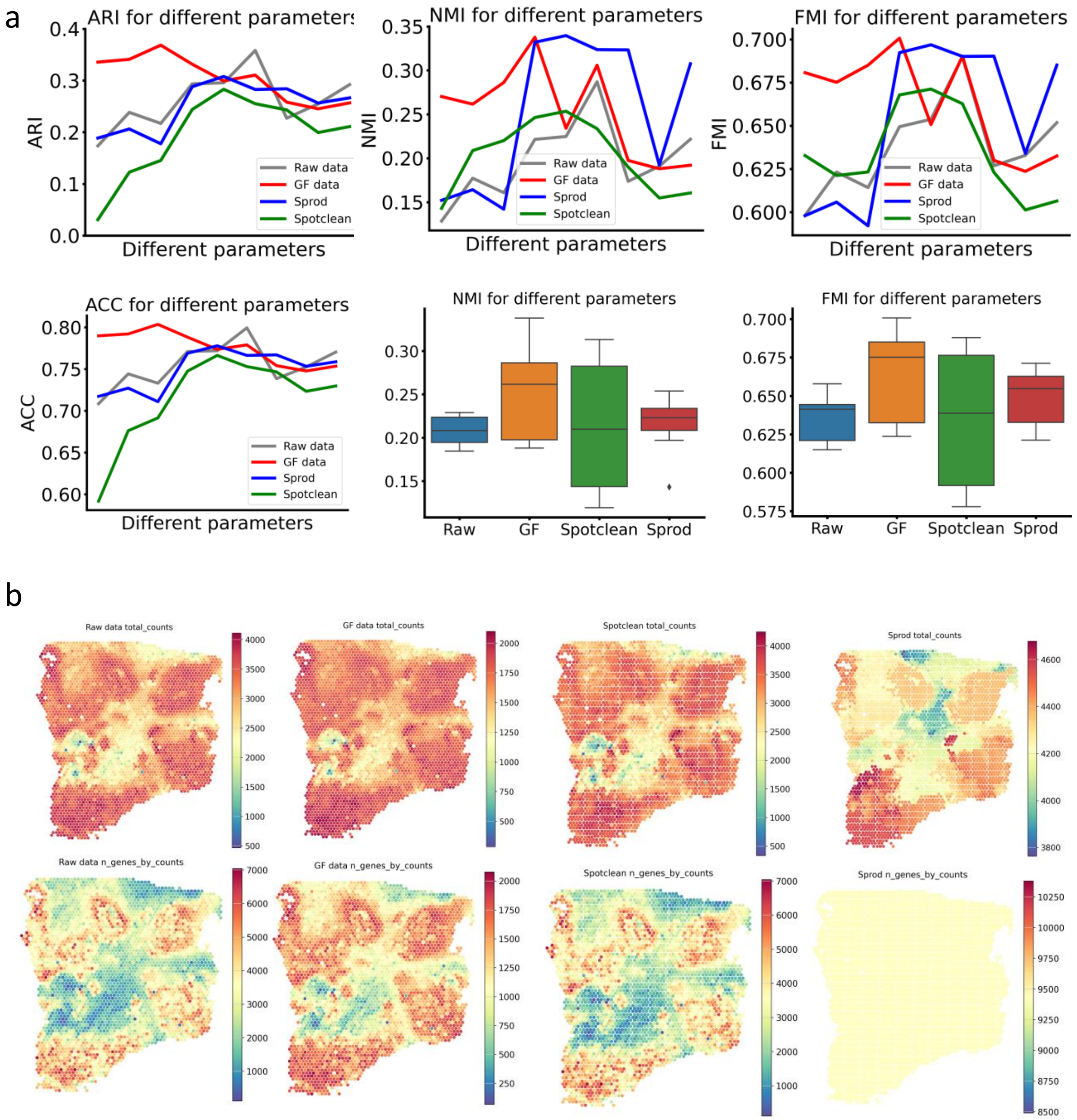

**Extended Data Fig.6**

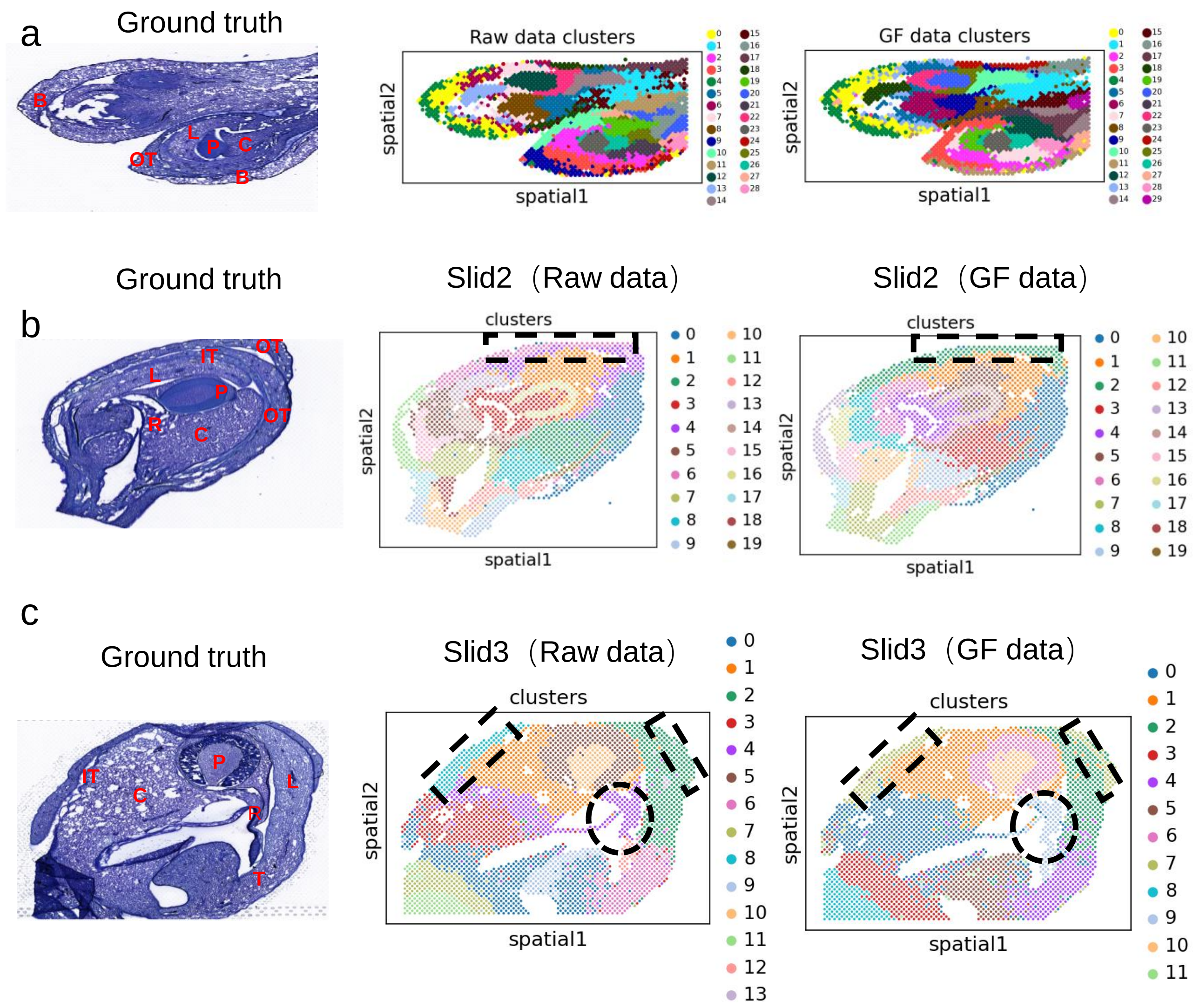


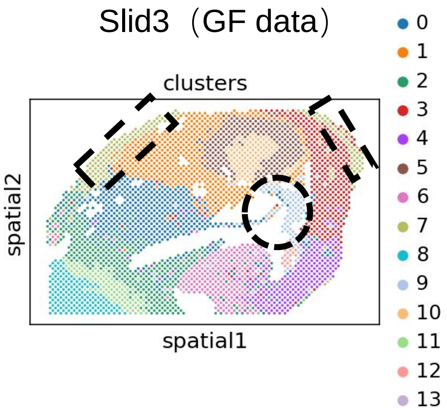


d

| abbreviation | tissue | abbreviation | tissue |
|---|---|---|---|
| B | Bract | OT | OuterTepal |
| P | Pollinium | IT | Inner Tepal |
| FM | Floral Meristem | C | Column |
| FP | Floral Primordium | L | Lip |
| R | Rostellum | T | Tegula |

**Extended Data Fig.7**

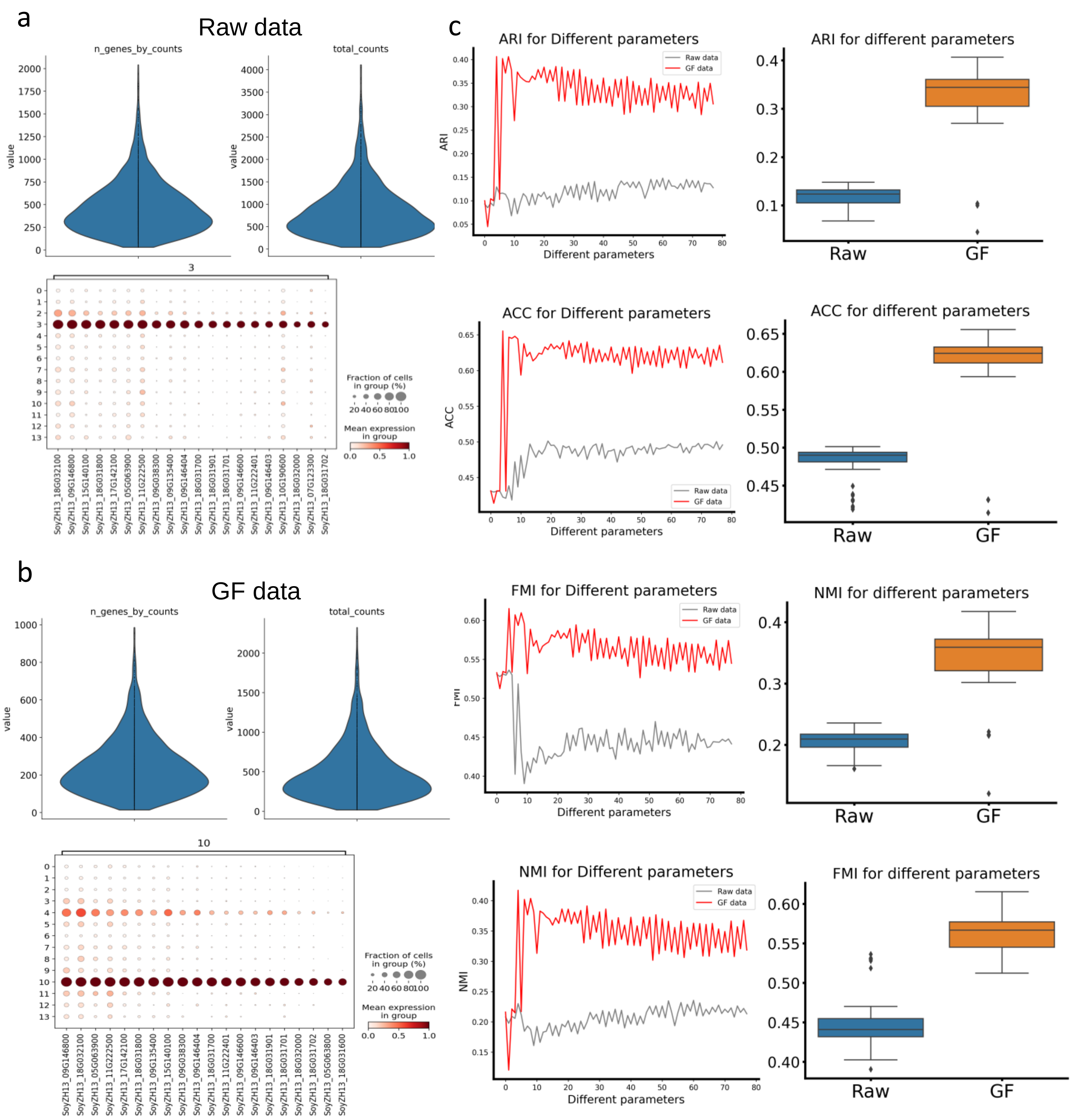

**Extended Data Fig.8**

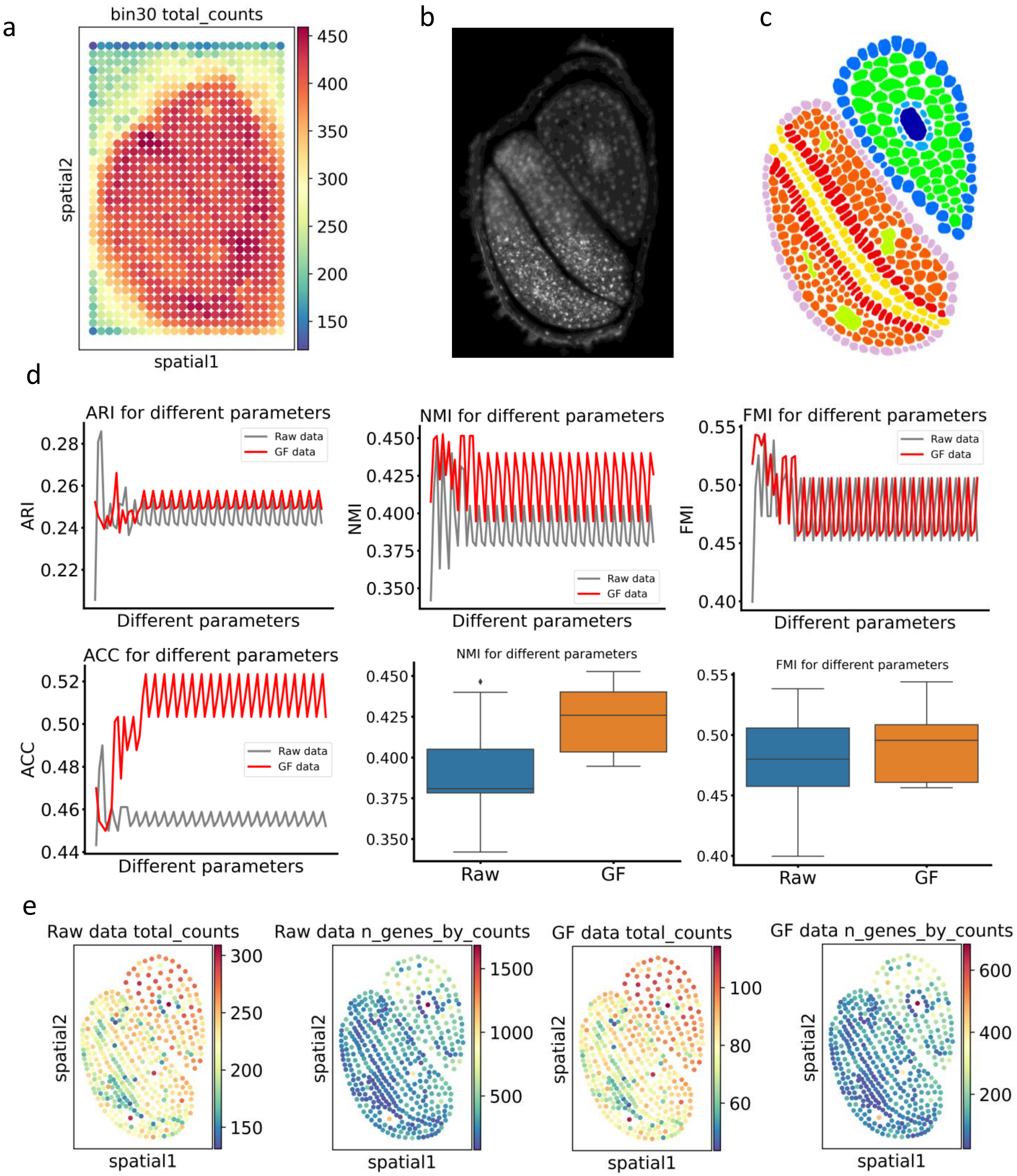

**Extended Data Fig.9**

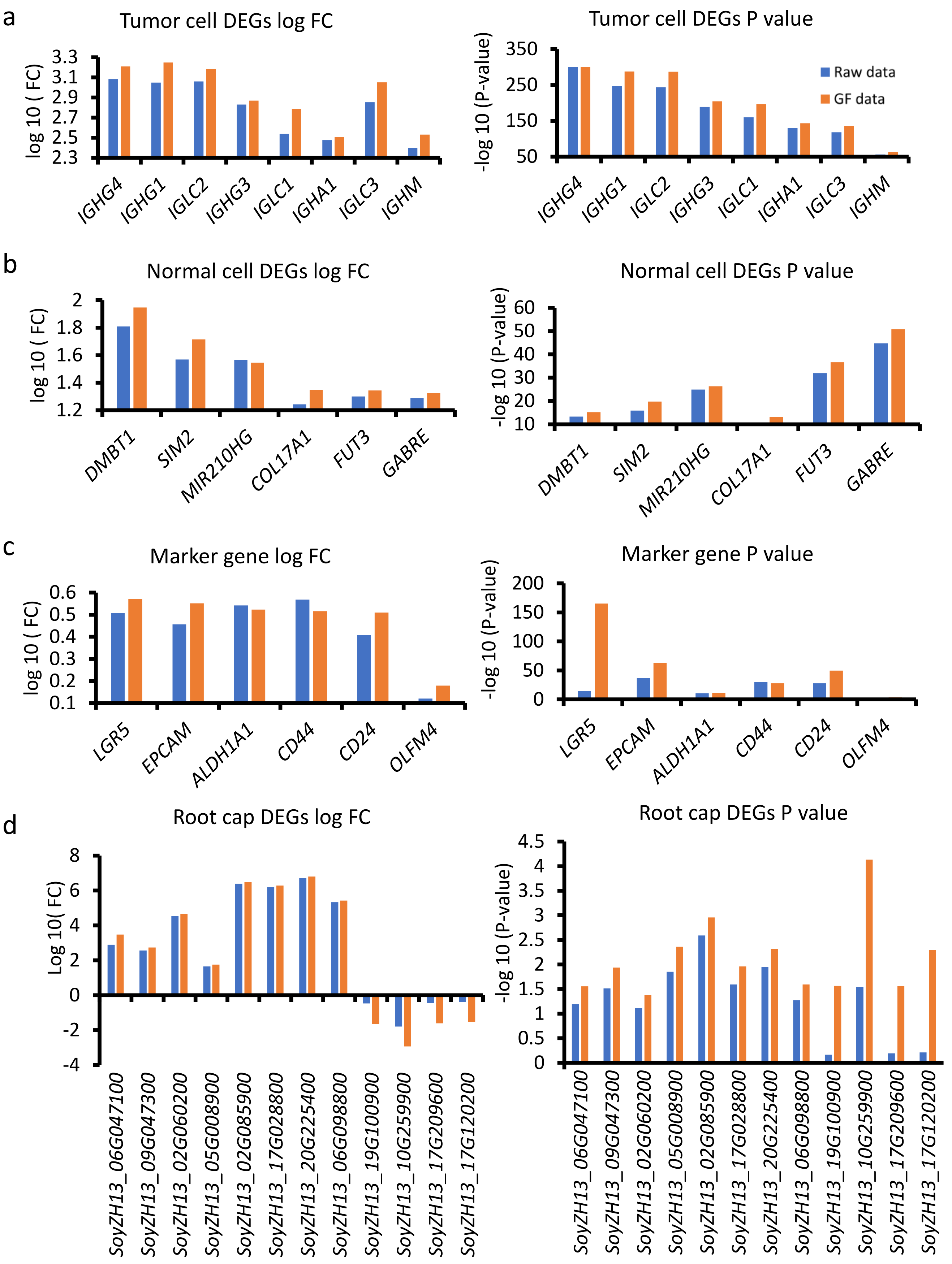

**Extended Data Fig.10**

a

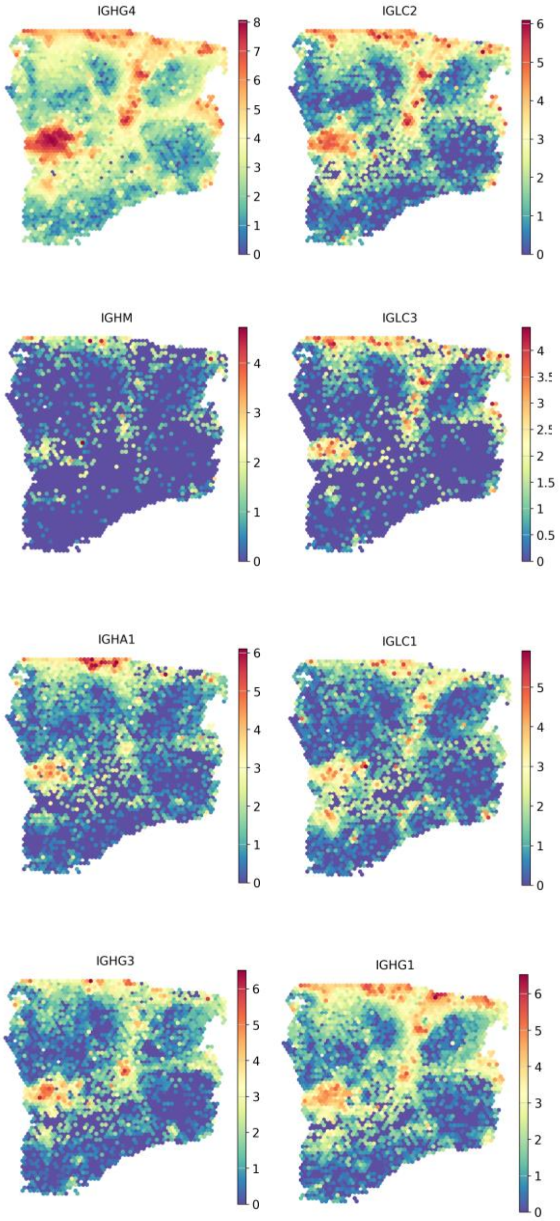


b

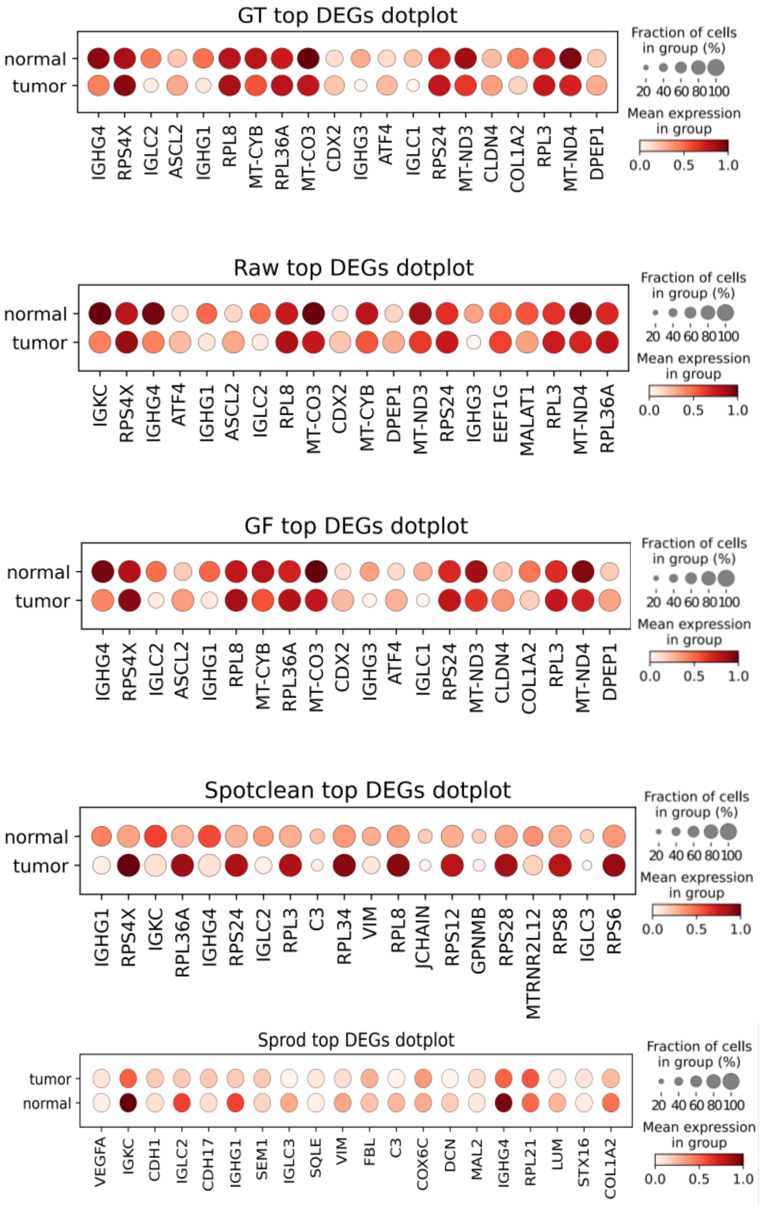
