## Supplementary material for "Optimal Transport Method-Based Gene Filter (GF) Denoising Algorithm for Enhancing Spatially Resolved Transcriptomics Data": Supplement figures legend.pdf

**Extended Data Fig. 1 | Randomly uniform noise added to the soybean SRT data and diffusion phenomena in other 10X Visium.**

**a,** Heat map of the sum distribution of UMI counts per cell in the raw SRT data and the noise added SRT data in figure 2e, and the distribution map of some noise genes.

**b,** Spatial expression distribution of the marker genes *PAXXGOG9220* and *PAXXG024270* of the bract and tepal tissue in the orchid slid1.

**c,** Spatial expression distribution of the marker genes *PAXXG048200* and *PAXXG086720* of the tepal tissue in the orchid slid2 data and the marker genes *PAXXG057040* and *PAXXG165320* of the meristem tissue in the orchid slid3 data. Spatial expression distribution of the marker genes *MOBP* and *GFAP* of the tepal tissue in the colorectal cancer data (slid: LIBD_151507).

**Extended Data Fig. 2 | Generation of Gaussian distributions with different degrees of clustering at random.**

The center point of all Gaussian distributions is (0,0), and four sets of different points are set, which are 500 spots, 1000 spots, 2000 spots, and 5000 spots. Set four sets of different covariance matrices to generate Gaussian distributions with different degrees of aggregation, in order [[2,1], [1,2]], [[5,1], [1,5]], [[10, 1], [1,10]], [[20,1], [1,20]], using the covariance matrix to represent the clustering degree of the two-dimensional Gaussian distribution.

**Extended Data Fig. 3 | Validating GF can accurately assess the degree of gene dispersion and is more effective than Moran's I.**

**a,** 10,000 different two-dimensional spatial distributions with a specified number of points were used constructed by Gaussian distribution and used the covariance matrix of the Gaussian distribution to characterize the degree of aggregation of each Gaussian distribution. The abscissa is different two-dimensional spatial distribution, and the ordinate is the degree of gene diffusion calculated by GF.

**b,** Moran's I coefficient is used to calculate the diffusion coefficient of each Gaussian distribution. The abscissa represents different two-dimensional spatial distributions, and the ordinate is the degree of diffusion of genes calculated by Moran's I. In a and b, green in the figure means that each distribution has 500 spots, red means that each distribution has 1000 spots and yellow and blue means that each distribution has 2000 spots.

**c,** Construction of 10,000 different two-dimensional spatial distributions of the same aggregation degree using a Gaussian distribution. The x-axis represents the number of points for each two-dimensional spatial distribution, and the y-axis represents the degree of spread of each distribution calculated using GF.

**Extended Data Fig. 4 | Confusion matrix of 10X Visium colorectal cancer data clustering results.**

According to the clustering results of Raw, GF-denoised, SpotClean-denoised, and Sprod-denoised SRT data in Fig.4c, we, calculated the correct rate of all cell predictions and draw four confusion matrices.

**Extended Data Fig. 5 | 10X Visium colorectal cancer data Supplementary test results.**
**a,** In the clustering results in Figure 4b, the unsorted line graphs of ARI, ACC, FMI, and NMI values. The x-axis represents 9 different parameters by varying the cluster from 2 to 4 and the learning rate from 0.1, 0.01, to 0.001,and the y-axis represents the planting of different indicators) and boxplots of FMI, and NMI values.
**b,** Heat maps of total counts expression per cell and heat maps of the number of genes expressed in each gene for raw SRT data, GF-denoised SRT data, SpotClean-denoised SRT data, and Sprod-denoised SRT data.

**Extended Data Fig. 6 | Improved clustering on 10X Visium Orchid Slice 2 and Slice 3.**
**a,** Clustering results of raw SRT data and GF-denoised SRT data. Scanpy software under the parameter conditions of resolution=3.0 and pc=30.
**b,** H&E staining map of orchid slid2 data. The clustering results of the raw SRT data of orchid slid2 data and the GF-denoised SRT data, Scanpy under the same parameter conditions (resolution=1.6, pc=30).
**c,** H&E staining map of orchid slid3 data. The raw SRT data is clustered into 14 categories, and the GF-denoised SRT data is clustered into 12 categories. Clustering using Scanpy software under the same parameter conditions (resolution=1.0, pc=50). GF-denoised SRT data gather 14 clusters when the parameters (resolution=1.5, pc=50).
**d,** Statistical table of correspondence between abbreviations and original words in 10X Visium orchid organization.

**Extended Data Fig. 7 | Supplementary results of Stereo-seq soybean data.**
**a,** Violin diagram of total counts expression per cell and the number of genes expressed in each gene for raw SRT data and the bubble diagram of the top 20 DEGs in cluster 3 of the raw SRT data clustering results in Fig.5f
**b,** Violin diagram of total counts expression per cell and the number of genes expressed in each gene for raw SRT data and the bubble diagram of the top 20 DEGs in cluster 3 of the GF-denoised SRT data clustering results in Fig.5f.
**c,** The unsorted line graphs of ARI, ACC, FMI, and NMI values calculated by SpaGCN. The x-axis represents 51 different parameters varying the cluster from 3 to 19 and the learning rate from 0.1, 0.01, to 0.001, and the y-axis represents the planting of different indicators and boxplots of FMI, and NMI values.

**Extended Data Fig. 8 | Supplementary results of Stereo-seq Arabidopsis data.**
**a,** In the clustering results in Figure 6a, the unsorted line graphs of ARI, ACC, FMI, and NMI values. The x-axis represents 41 different parameters varying the cluster from 5 to 18 and the learning rate from 0.1, 0.01, to 0.001, and the y-axis represents the planting of different indicators, and boxplots of FMI, and NMI values.
**b,** Heat maps of total counts expression per cell and heat maps of the number of genes expressed in each gene for raw SRT data and GF-denoised SRT data.
**c,** The Heat map of Stereo-seq Arabidopsis data bin size 30 expression, FB staining image, and cell

segmentation diagram (Ground truth).

**Extended Data Fig. 9 | Log fold change and -log 10 (P value) of DEGs identified by raw data and GF data.**

**a,** Log fold change (log FC) and the -log 10 (P value) of eight DEGs (immunoglobulin marker genes) reported in colorectal cancer tumor cells in the raw and GF SRT data clustering results in Fig.4c. Set to 300 if the -log 10 (P-value) is greater than 300.

**b,** Six DEGs were highly expressed in normal cells, and Log fold change (log FC) and -log 10 (P value) were calculated in the clustering results in Fig.4c using raw SRT data and GF-denoised SRT data respectively.

**c,** Log fold change and -log 10 (P value) in the clustering results in Fig.4c were calculated for the marker genes of 6 tumor cells reported in the cell marker database, in the raw SRT data, and in the GF denoised SRT data.

**d,** Log fold change and -log 10 (P value) for the same DEGs in the root cap tissue calculated by the t-test method in the clustering results in Fig.5f. Cluster 3 in raw SRT data clustering result and Cluster 10 in GF SRT data clustering result.

**Extended Data Fig. 10 | The bubble plot of DEGs and 8 known tumor cell marker genes in the clustering results of Figure 4c.**

**a,** Expression bubble chart of top 10 DEGs identified by tumor cells and normal cells, using t-test method to identify DEGs, across from Ground truth SRT data, raw SRT data, GF-denoised SRT data, SpotClean-denoised SRT data.

**b,** The Expression heat map of immunoglobulin marker genes in 8 reported 10X Visium colorectal cancer tumor data.
