## Supplementary material for "Optimal Transport Method-Based Gene Filter (GF) Denoising Algorithm for Enhancing Spatially Resolved Transcriptomics Data": Supplementary tables legend.pdf

**Supplementary table1：** Clustering result indicators of various organizations treated differently

**Supplementary table2：** The soybean Stereo-seq adds 5000 noise genes data to calculate the contingency table of each cell prediction result and Ground truth annotation result with SpaGCN

**Supplementary table3：** The soybean Stereo-seq raw data calculate the contingency table of each cell prediction result and Ground truth annotation result with SpaGCN

**Supplementary table4：** Parameters traversed in several processing flows

**Supplementary table5：** GF results of different Two-dimensional Gaussian distribution dataset

**Supplementary table6：** GF results of different dataset

**Supplementary table7：** The result of traversal when using SpaGCN to cluster colorectal cancer 10X Visium data

**Supplementary table8：** The result of traversal when using Scanpy to cluster soybean root data (78 groups)

**Supplementary table9：** The soybean Stereo-seq raw data calculates the contingency table of each cell prediction result and Ground truth annotation result with Scanpy(resolution=0.65)

**Supplementary table10：** The soybean Stereo-seq raw data calculates the contingency table of each cell prediction result and Ground truth annotation result with Scanpy(resolution=1.30)

**Supplementary table11：** The soybean Stereo-seq GF data calculate the contingency table of each cell prediction result and Ground truth annotation result with Scanpy(resolution=1.30)

**Supplementary table12：** The result of traversal when using SpaGCN to cluster Arabidopsis root data (41 groups)

**Supplementary table13：** The Arabidopsis Stereo-seq raw data calculates the contingency table of each cell prediction result and Ground truth annotation result

**Supplementary table14：** The Arabidopsis Stereo-seq GF data calculates the contingency table of each cell prediction result and Ground truth annotation result
